# Generative semantic multiplexing (SemaPlex) for accessible and scalable multiplexed fluorescence imaging

**DOI:** 10.1101/2025.09.15.676186

**Authors:** Ihuan Gunawan, Moumitha Dey, Daniel P Neumann, Felix V Kohane, Yaqing He, Erik Meijering, John G Lock

## Abstract

Multiplexed fluorescence imaging enhances spatially-resolved interrogation of complex, multi-molecular cell processes that are insufficiently sampled using standard 4-5 plex imaging. To improve accessibility and scalability for multiplexed imaging, we demonstrate generative ‘Semantic Multiplexing’ (SemaPlex); a simple experimental and deep learning strategy for amplifying marker plexity several-fold by semantically unmixing multiple markers combined per imaging channel. We first characterise key determinants of SemaPlex performance, achieving precise computational multiplexing of 2-to-8 markers synthetically mixed in one channel, facilitating enhanced cell phenotype classification. We then demonstrate practical SemaPlex application, acquiring 10 markers over 4 channels (3*3-plex+1) to efficiently emulate real multiplexed labelling. This permitted accurate reconstruction of quantitative single-cell phenotypic manifolds delineating cell-cycle and mitotic dynamics, with internally validated error-detection. Finally, we exemplify use of ‘semantic guides’; additional input channels that significantly enhance multiplexing fidelity. SemaPlex makes multiple-fold increases in fluorescence imaging-plexity accessible, scalable and customisable; democratising multiplexed imaging-based interrogation of complex cell biology.

## Introduction

Fluorescence microscopy has long enabled single-cell data analyses that access subcellular resolution not captured by most single-cell omics^1^. Imaging-based systems biology approaches^2^ have been leveraged in various settings, especially in high-throughput applications detailing cellular phenotypes and their molecular underpinnings^3,4^. Yet standard fluorescence imaging techniques detect just 3 to 5 molecular markers per sample (3- to 5-plex) due to limitations including fluorophore spectral-crosstalk^5^ and antibody cross-reactivity. This low-plexity profoundly constrains detailed interrogation of complex, multi-molecular cell processes. To enrich fluorescence imaging-based data, experimental techniques have been developed to permit multiplexed imaging of (typically) 10-30+ markers per sample. Methods such as 4i^6^ and CycIF^7^ have become the gold-standard of open-source experimental multiplexing and have, for example, enabled recognition of links between subcellular multi-molecular organisation and cell states^8^, understanding of how cell states modulate single-cell perception of incident stimuli^9^, and mapping of multi-molecular cell state transitions in cancer tissue^10^. However, the relatively high experimental burden associated with such methods constrains their widespread application, particularly in the contexts of high-throughput studies such as phenotypic screening where cyclic labelling / imaging requirements become prohibitive. Alternatively, spectral unmixing methods can also achieve increased marker plexity (typically 7-10 per sample) while maintaining greater experimental scalability. Combinations of spectral unmixing and CycIF have achieved high-plexity imaging across multiple labelling and spectral imaging cycles^11^, yet such approaches may be even more restricted in application due to combining the demands of cyclic imaging with requirements for specialised spectral imaging hardware. Widely employed phenotyping approaches like Cell Painting^12^ achieve modest multiplexing at high-throughput by combining molecular markers/dyes to label multiple subcellular structures in the same channel. While broadening assay sensitivity to detect phenotypic effects across a wider array of subcellular structures, the specific biological interpretability of such mixed marker channels is reduced and the assay does not readily permit monitoring of bespoke molecular targets^13,14^. To limit the confounding effects of in-channel marker mixing and to enable some bespoke marker inclusion, Coburg *et al.* presented Cell Painting PLUS, which employs sequential imaging and elution to separate acquisition of each dye^14^; enhancing specific biological interpretability while permitting some bespoke molecular labelling. This effort emphasises the value to the field of increasing plexity by even 1-2 additional markers, yet the requirement for multiple-cycle iterative labelling in the applied (e.g. screening) context imposes a significant experimental burden that may preclude widespread use.

Beyond experimental multiplexing methods, advances in deep-learning now permit predictive or ‘virtual’ marker labelling, leveraging model inputs including label-free images and/or other fluorescent markers^15,16^. Indeed, multiple input channels can enhance predicted label fidelity while also guiding anchored integration of numerous predicted markers, achieving robust computational multiplexing^16^. Yet although recent studies show that computational multiplexing using virtual markers can significantly enhance downstream phenotypic and molecular analyses of single-cell populations^16^, the fidelity between virtual and ground-truth images remains of an intermediate level^15,17,16^ (∼0.5-0.7 Pearson’s Correlation Coefficient across pixels). Ultimately, this moderate fidelity may limit individual virtual marker interpretability and thus the overall applicability of these essentially predictive virtual labelling methods. Computational approaches are also now emerging to separate multiple fluorescently labelled markers (or structures) acquired within a given imaging channel, moving beyond the predictions of virtual labelling to a form of deconvolution based on real signals that may facilitate substantially higher ground-truth fidelity. Using a feature extraction-based approach, Ben-Uri *et al*. showed that a machine learning model can discriminate between the segmented, quantified morphological characteristics of two markers (Golgi, mitochondria) mixed in a single channel during live imaging, permitting their parsing and independent analysis^18^. Zhanghao *et al.* split non-specific dye labels spanning multiple subcellular structures into distinct images of specific organelles-of-interest (whose labelling comprised a subset of the total non-specific labelling) based on training deep learning models using parallel ground-truth organelle labelling^19^. However, such nonspecific dyes predetermine the set of cellular structures labelled, precluding selection of custom markers/targets. By contrast, careful design and labelling of bespoke marker combinations in sequential imaging cycles can permit a form of spectral signal-barcoding allowing high-fidelity generative reconstruction of customisable molecular marker panels^20^. Enabling more cycle-efficient multiplexing than methods such as 4i and CycIF, this ‘CombPlex’ approach demands labelling of most markers in multiple independent channels across multiple acquisition cycles, necessitating a complex experimental protocol that may limit utilisation of this otherwise powerful strategy.

Given this landscape of existing methods, we identified the need for a more accessible and scalable approach to multiplexed imaging. We thus present Semantic Multiplexing (SemaPlex), an integrated experimental and deep learning strategy permitting each available imaging channel to host multiple (2 to 8 shown herein) specific molecular markers bearing equivalent fluorescent tags. Semantic (as opposed to spectral) unmixing then leverages differences in the spatial patterning of marker signals (even when spatially overlapping), learned by deep convolutional neural networks, to produce high-fidelity individual reconstructions of each marker present in a given channel. The SemaPlex strategy incorporates a simple, compact experimental design requiring only standard (immuno)fluorescence labelling reagents and imaging hardware for initial deep learning model-training (schematised for 2-plex per channel in **Figure 1A, B**). SemaPlex application (**Figure 1C, D**) is then highly efficient and minimally complex, requiring only a ‘single-shot’ mixed marker acquisition. This makes SemaPlex a highly scalable (e.g. for high-throughput screening) strategy by which to amplify bespoke marker plexity several-fold per imaging channel. We ultimately show that SemaPlex achieves fidelity to ground-truth labelling exceeding that of virtual labelling methods, instead being comparable to repeat experimental labelling and imaging of same target marker – a realistic upper bound on possible performance. To further enhance SemaPlex, we have implemented a powerful error-detection method to detect and (with tuneable thresholding) remove sub-optimal marker reconstructions at single-cell resolution (**Figure 1E**). We also present ‘semantic guidance’ as an extension of SemaPlex that leverages additional available imaging channels (e.g. nuclear or label-free images) to enhance and correct the unmixing of challenging marker combinations (**Figure 1F**). Semantic guide images provide additional biological context to generative SemaPlex models, significantly increasing the fidelity of reconstructed target images and providing extensive scope for bespoke optimisation of the SemaPlex approach.

**Figure 1.**
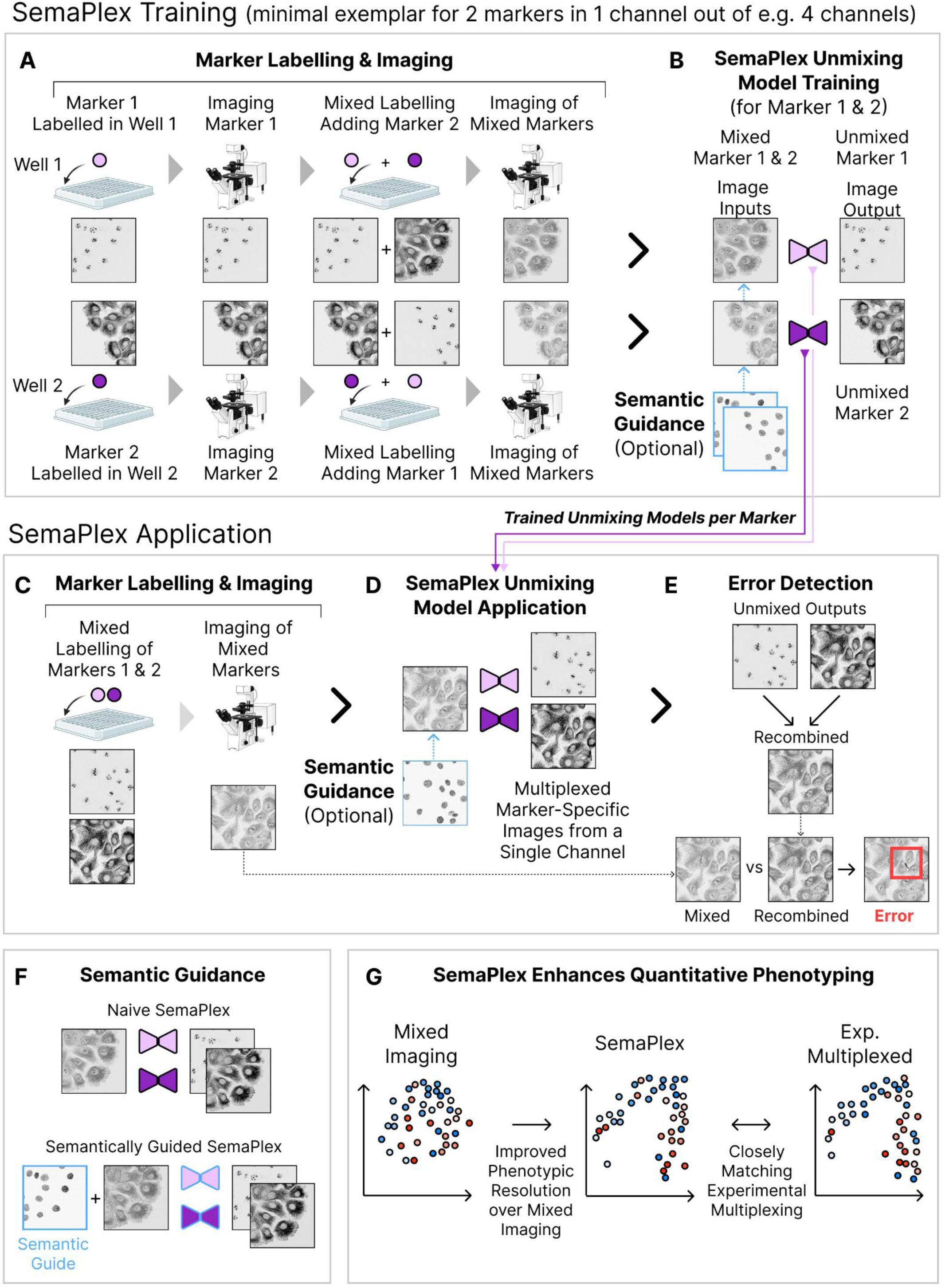
Schematic overview of SemaPlex training and application workflow. **(A)** SemaPlex training reflects a compact experimental design, wherein the number of cell populations (e.g. wells) required for deep learning model training is equal to the maximum number of markers mixed per channel. For simplicity, a minimal marker labelling and acquisition schema is shown here for 2-plex mixed marker acquisition in 1 channel. Using standard (immuno)fluorescence labelling and imaging, each marker to be unmixed per channel is first separately labelled and imaged within a different well (2 wells needed for 2-plex unmixing). The remaining marker(s) to be unmixed per channel are then added to respective wells, enabling mixed marker image acquisition. **(B)** SemaPlex deep learning models (one per mixed marker) are then trained to reconstruct each individual marker (model targets) from the mixed marker images (model inputs), producing semantically multiplexed datasets. SemaPlex models can optionally take additional input(s) for semantic guidance, explained further below in panel F. **(C)** Post-training, SemaPlex applications are minimally complex and highly scalable, with mixed marker images able to be generated at high-throughput via labelling and ‘single-shot’ acquisition of all mixed markers at once. **(D)** Pre-trained SemaPlex models then precisely reconstruct images of each specific target marker from these (unseen) mixed marker images. Again, SemaPlex models can optionally take additional input(s) for semantic guidance, explained further below. **(E)** Internally validated error correction is integrated within the SemaPlex workflow by recombining unmixed markers and comparing the recombined and original mixed marker images, revealing erroneous single-cell reconstructions that can be filtered from downstream data. **(F)** SemaPlex performance can be further enhanced by including additional ‘semantic guide’ channels as model inputs, providing enhanced context to guide marker unmixing. **(G) Left:** Expanding the schema outlined above, SemaPlex can be applied across multiple channels, with multiple mixed markers per channel, to produce high plexity datasets wherein the biological trends of each individual marker can be extracted, rather than being convoluted within a mixture of markers. **Right:** SemaPlex also enhances quantitative single-cell phenotyping, as captured via multivariate manifold embedding, wherein mixed image data yields poorly resolved information on single-cell heterogeneity. By contrast, SemaPlex-unmixed images of each marker collectively yield more nuanced single-cell phenotype resolution, producing unsupervised data manifolds closely recapitulating ground-truth single-cell biology captured from the same cells via experimentally multiplexed data.

Though SemaPlex has been independently conceived and developed, we note that MicroSplit^21^, recently proposed by Ashesh *et al.,* applies similar unmixing concepts with a focus on minimising imaging illumination dose; vital in the context of live imaging where phototoxic effects require careful management^22^. Chen *et al.* similarly demonstrate capacity to unmix markers within a single channel during live imaging^23^. While Ashesh *et al.* and Chen *et al.* predominantly present high-fidelity examples of synthetic / simulated marker multiplexing in a single channel (up to 4 markers or up to 2 markers, respectively), we present both synthetic (up to 8 markers/channel) and real experimentally mixed marker reconstruction (up to 3 markers with the same fluorophore in a single acquisition channel) across multiple channels. In our initial characterisation of SemaPlex utility using synthetically mixed markers, we mapped the impacts on performance of factors including relative marker intensity within a channel, marker spatial (co)distribution, and marker number per channel. We then demonstrate that SemaPlex enhances cell phenotype classification accuracy beyond that achieved by using the mixed marker (input) channels. Building on these ‘simulated’ applications, we next applied SemaPlex to generate real 10-plex data including DAPI alone (one channel) plus 3 markers per channel across 3 channels. With markers sampling cell cycle and mitosis biology, we applied single-cell quantitative analyses and unsupervised manifold learning to infer cell cycle distributions and to stage mitotic phases, comparing SemaPlex-derived multiplexed data to ground-truth experimentally multiplexed (CycIF) data (same markers, same cells) acquired after imaging of SemaPlex input data (**Figure 1G**). This confirmed that data multiplexed via SemaPlex delineates phenotypic diversity with a resolution mirroring that achieved via far more demanding experimentally multiplexed labelling.

Taken together, SemaPlex can amplify fluorescence imaging marker-plexity by several fold, whilst also being compatible with other (e.g. cyclic and/or spectral) multiplexing techniques. Achieving high-fidelity image-and biological process-reconstruction, SemaPlex requires only standard (immuno)fluorescence reagents and hardware, making it highly accessible. SemaPlex is also scalable in molecular plexity as well as being compatible with high-throughput (e.g. screening) applications, whilst maintaining marker selection flexibility and marker-specific interpretability. Thus, SemaPlex opens the door to more bespoke and informative cell phenotyping via multi-molecular imaging assays, at scale; a substantial advance for imaging-based research capabilities.

## Results

### SemaPlex Unmixing Validation in Image Datasets with Simulated Mixing

SemaPlex employs deep learning to computationally isolate mixed markers imaged in a single channel. Intuitively, using mixed markers as a deep learning input explicitly includes signals necessary to unmix individual target markers, allowing high-fidelity multiplexed reconstruction. Yet, experimental factors including selection of marker combinations and relative marker labelling intensity could impact multiplexing performance. Accordingly, we first characterised SemaPlex unmixing performance using ‘synthetically’ mixed data, allowing us to control and thereby evaluate the influence of these factors, to test many marker combinations, and to assess various numbers of markers (synthetically) mixed in a single channel. To enable this extensive analysis and validation, we utilised a CycIF-based, experimentally multiplexed dataset from HCC38 breast cancer cells treated with transforming growth factor-beta one (TGF-β1) for 20 minutes, 1 hour, or 3 hours to induce the preliminary phases of epithelial-to-mesenchymal transition (EMT) and thereby modestly diversify cell phenotype heterogeneity. Direct immunofluorescence labelling of molecular markers then included: α-tubulin; β-catenin; cytokeratin; DNA (DAPI); fibrillarin; phospho-SMAD2/3 (p-SMAD2/3), CD44, and vimentin (**Supplementary Table 1**). Crucially, we note that experimentally multiplexed data was used purely for exploratory testing and validation of SemaPlex unmixing. Neither training nor application of SemaPlex unmixing requires multiplexed data. SemaPlex can be readily performed with standard 4-plex fluorescence imaging data.

All synthetic (computational) mixing of markers for exploratory analyses was performed by calculating pixel-wise mean intensity from all channels being mixed. SemaPlex unmixing was then performed on the mixed images using a deep convolutional neural network with U-Net^24^-like architecture (**Methods**), trained to semantically unmix and thereby reconstruct the individual molecular marker ‘targets’ from within the mixed marker image, as depicted in **Figure 1B**. Unless otherwise indicated, all analyses herein employ 5-fold cross-validation, with unmixed marker image fidelity (to the matched ground-truth marker image) quantified using pixel-level Pearson’s correlation coefficient^25^ (Pixel-PCC); recently shown to best reflect further downstream analysis-utility relative to other image quality metrics^26^.

### High Fidelity Semantic Multiplexing from Simulated Mixed Data

We first tested the reconstruction of individual marker images from mixed images combining various marker pairs, exploring the impact of marker spatial overlap on reconstruction performance. Across all pairs, we found that markers could be accurately recreated in individual channels with high visual fidelity (**Figure 2A**). Quantitatively, all tested marker combinations achieved Pixel-PCC scores above 0.85, with several approaching perfect reconstruction (Pixel-PCC ≐ 1) (**Figure 2B**). Unsurprisingly, markers with low spatial overlap and/or distinctive local pixel patterning, such as pSMAD2/3 and Vimentin, or DNA and fibrillarin, generally achieved higher Pixel-PCC values (∼0.95 or higher). In contrast, markers with high spatial overlap and similar patterning, like CD44 and β-catenin, achieved slightly lower-fidelity reconstruction (∼0.9). This offers a guide to inform SemaPlex panel design, as exemplified in our subsequent application. While our results demonstrate the efficacy of semantic unmixing in ideal labelling conditions, fluorescent markers may in practice have significantly different signal strengths that may unbalance their presentation when mixed in a single channel. We simulated this scenario by attenuating the signal of one marker while keeping the other at full strength before mixing the channels (**Figure 2C**). Despite a significant drop in performance with larger attenuation (and a corresponding though diminishing improvement in the marker with dominant intensity), reconstruction performance remained very high across most conditions (**Figure 2D**). In fact, Pixel-PCC scores only fell below 0.75 when the attenuated marker was presented at 10% strength, establishing a highly imbalanced, order-of-magnitude difference in signal between the synthetically mixed marker pair.

**Figure 2.**
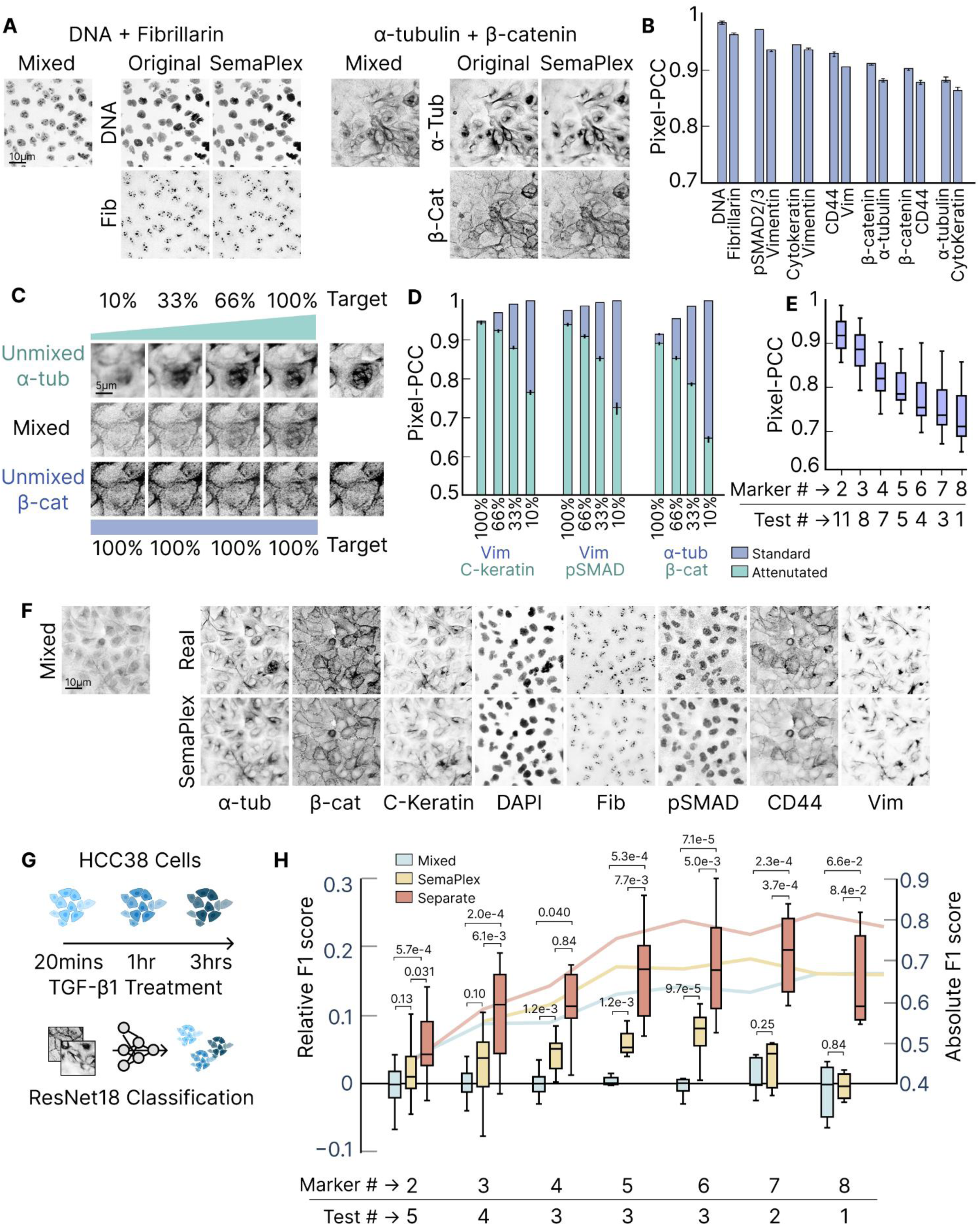
Characterisation of SemaPlex performance with synthetically mixed labelling. **(A)** Examples of marker unmixing. Mixed images generated by taking the pixel-wise mean of separate marker images were used as deep learning model-input, with the individual markers as targets for model training. Trained models were applied to unseen mixed images to demonstrate visual fidelity for each marker (upper and lower rows as respectively titled). **(B)** Pixel-PCC scores across 5 cross-validated folds for various unmixed marker pairs. **(C)** Example cropped image field showing visual unmixing performance with different levels of marker intensity-imbalance. Target column in top and bottom rows show real experimental labelling. Remaining columns show unmixed images when target markers are mixed (middle row) at intensity percentages as indicated. **(D)** Pixel-PCC scores across 5 cross-validated folds quantifying unmixing fidelity for imbalanced mixed images as exemplified in C, including alternative marker pairs as indicated (X axis). **(E)** Pixel-PCC scores across 5 cross-validated folds per marker for separation of increasing marker plexity (Marker #; 2 to 8) mixed per image enabling higher SemaPlex-mediated multiplexing. Boxes show average performance for each unmixed marker across 5 cross-validated folds. Number of marker combinations tested per plexity (Test #) is n=11, 8, 7, 5, 4, 3, 1 respectively for 2 to 8-plex combinations. Marker combinations used detailed in **Supplementary Table 2**. **(F)** Example image containing 8 mixed markers (left), the ground-truth individual markers (right, upper row) compared to the unmixed markers (right, lower row). **(G)** Experimental design for downstream evaluation of unmixing efficacy in cell phenotype classification. HCC38 cells underwent TGF-β1 treatment for three different durations to induce subtle phenotypic changes associated with early-stage epithelial-mesenchymal transition (EMT). ResNet18 (deep learning model-based) classification of treatment condition (time) was then performed using images of various compositions (detailed in H). (**H**) Treatment condition-classification, detailed in G, leveraged image mixtures of increasing marker plexity (Marker #; marker combinations used detailed in **Supplementary Table 2**). Number of marker combinations tested per plexity (Test #) is n=5, 4, 3, 3, 3, 2, 1 respectively for 2 to 8 -plex. Markers were either: mixed in a single channel or; Semantically Multiplexed (SemaPlex) via deep learning-enabled unmixing from the same single mixed channel, or; used in their ground-truth (experimentally multiplexed via CycIF) form with one marker per image channel. Classification performance was quantified as median F1-score over the last 50 epochs (from a total of 300 epochs) of ResNet18 optimisation, across 3-folds cross-validated by holding out one well per condition. Boxes show F1-Score performance relative to the mixed marker images (left y-axis), centred around 0 by subtracting the median mixed score (across the folds) for each combination of markers. Median absolute F1 score shown as a line for each condition (right y-axis). Error bars/box plots in all panels show lower and upper quartiles. Significance tests reflect Welsh’s two-sided t-test. Scale bars = 10μm.

We next tested unmixing and reconstruction for larger numbers of markers per mixed channel, to assess the capacity of SemaPlex to generate higher plexity datasets. We tested a total of 39 marker combinations mixing from two to eight markers in a single channel (**Figure 2E**, **Supplementary Table 2**). As expected, reconstructed image fidelity gradually declined with mixing of greater marker numbers, though even with 8 co-mixed markers, Pixel-PCC values concentrated between 0.7 and 0.8 (image examples in **Figure 2F**). Overall, our results show that high-fidelity semantic multiplexing is effective even for large numbers of (mixed) markers, and also where marker signals are up to 10-fold imbalanced.

### Unmixed Markers Improve Phenotyping Capacity

While there is clear value for biological interpretation in having distinct marker labelling, which SemaPlex enables, the extent to which biological information may be lost by ‘overloading’ marker labelling in a single channel remains unclear. Likewise, it remains to be seen whether SemaPlex can recover obscured/lost information from mixed marker channels to standards achieved through real experimental multiplexing. To answer these questions, we next performed quantitative classification (using the ResNet18 deep learning model) to discern subtle cell phenotypes emerging during early timepoints (20 min, 1h, 3h) of TGF-β1-treatment, which is a canonical driver of epithelial-to-mesenchymal transition (EMT)^27,28^ (**Figure 2G**). This relatively challenging task was selected to provide substantial scope for different levels of classification performance based on widely varied image data inputs. Specifically, we tested classification performance using various marker combinations of from 2 to 8 markers (combinations as per **Supplementary Table 2**) that were either: synthetically mixed, or; unmixed after synthetic mixing, or; in their original experimentally multiplexed form as separate channels (**Figure 2H**). In general, additional markers increased classification performance irrespective of the configuration of markers (mixed, unmixed, or separate). Classification using markers in separate channels increasingly outperformed the corresponding mixed inputs as marker numbers increased, implying information loss with increasingly overlapping (mixed) markers. Notably, unmixed channels generally improved classification outcomes over the mixed channels that were used as model-inputs though the original separate channels showed maximal performance. SemaPlex-mediated improvements were particularly significant for intermediate marker numbers per channel (4-6). At 8 mixed markers, SemaPlex performance reverts to match the mixed condition, likely due to limitations in recovering highly overlapping signals. These results ultimately suggest that overloading of markers into a single channel obscures phenotypic information, especially at 3 or more markers per channel, but that SemaPlex-enabled unmixing and multiplexing permits significant recovery or resurfacing of this convolved biological information, particularly in the range of 4-6 markers per channel. SemaPlex can thus substantially enhance capacity to discern subtle cell phenotypes.

### Experimental Application of Semantic Multiplexing

Having validated our approach on synthetically mixed images, we next demonstrate a real-world implementation of SemaPlex with experimentally labelled data. We showcase practical end-to-end application of the SemaPlex experimental labelling / imaging design and of SemaPlex model training and inference to achieve multiplexed (10-plex) labelling based only on standard four-channel fluorescence imaging. In addition to showing strong performance visually and at the fundamental level of pixel-wise PCC correlations, we confirmed via quantitative single-cell phenotyping that semantically multiplexed markers recapitulate higher-order biological insights at similar resolution to that afforded by experimental multiplexing, which we performed via CycIF after acquiring the data necessary for SemaPlex. We again stress that this experimentally multiplexed CycIF data was acquired here for comparative purposes only and is not required for SemaPlex implementation.

To exemplify the utility of SemaPlex for biologically informative single-cell phenotyping using bespoke molecular markers, we performed a representative downstream single-cell quantitative analysis of, in this case, cell cycle phenotypes and dynamics; thereby tackling a highly multi-molecular cell process involving coordinated changes in protein expression and localization, organelle organisation, and cellular morphology^29,30^. Specifically, we present concurrent measurement of various organelle / structural and cell cycle-related markers (**Supplementary Table 1**) including: DAPI (DNA); fibrillarin (Fib; nucleoli); β-tubulin (β-tub; microtubules); p21 (suppressed proliferation); ki67 (proliferation); Lamin A/C (Lamin; nuclear envelope); Cav-1 (Cav1; caveoli); PCNA (DNA synthesis); ZO-1 (ZO1; cell-cell tight junctions), and; SDHA (mitochondria). Acquiring 10 markers in total, we chose to label DAPI (excited at 405 nm) in its own channel, and to group remaining markers into mixtures of three markers per channel across each of three remaining channels (β-tub + Fib + p21 excited at 488 nm; Cav1 + ki67 + Lamin excited at 555 nm; PCNA + SDHA + ZO1 excited at 647 nm; **Figure 3A**). For each 3-plex mixed marker channel, markers were chosen based on having relatively distinctive spatial localizations and/or local pixel patterning, enabling robust unmixing performance.

**Figure 3.**
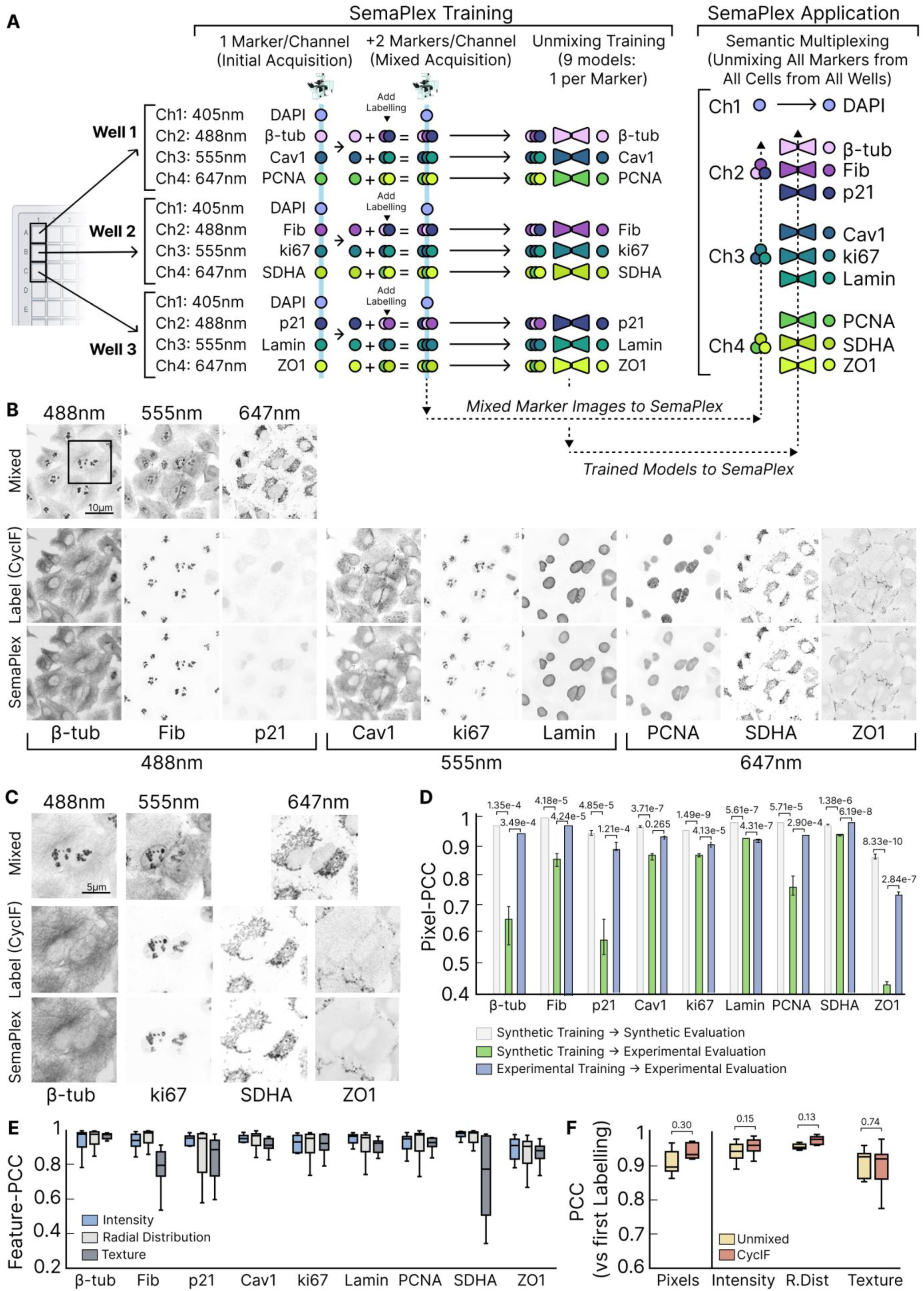
Evaluation of unmixing performance for experimentally mixed labelling. **(A)** Exemplar SemaPlex application schema starting with experimental labelling for 4-channel acquisition with DNA (alone) plus 3 channels each containing 3 markers, followed by unmixing to generate 10-plex Semantically Multiplexed data. Efficiently designed training data production here required only 3 cell populations (i.e. 3 wells); matching the number of markers in mixed channels. In each well, an alternate one of the three markers per channel was first individually labelled and imaged to acquire isolated images for that marker, creating training targets for all 9 markers to be unmixed. Next, the remaining two markers per unmixing channel were added and imaged to acquire mixed marker images comprising all three markers per channel; used as unmixing model-inputs. 9 SemaPlex unmixing models were thus trained; one each for the 9 distinct markers to be unmixed. All 9 trained models were then applied to the previously acquired mixed channels (see dashed arrow) to generate (in total) a 10-plex SemaPlex dataset spanning 4 standard fluorescence imaging channels. Not shown here for simplicity, we also generated ground-truth marker comparison data for all markers in all cells, using three CycIF cycles after mixed marker imaging to sequentially label all individual markers, enabling quantitative comparison of SemaPlex-*versus* experimentally multiplexed-images for all cells. **(B)** Example field showing marker unmixing in each channel. Mixed images per channel (top row) are compared to ground-truth cycIF images (middle row; acquired for comparison only) and to SemaPlex images (bottom row). Images contrast-scaled for visualisation purposes only. **(C)** Example crops of regions as marked in panel B. **(D)** Median Pixel-PCC (Pearson’s correlation coefficient) score across 5-cross validated folds for SemaPlex unmixed images compared to individually labelled and imaged, ground-truth marker targets. SemaPlex Models were either: trained using synthetic (computationally merged) images then evaluated on synthetic mixed marker images (grey bars) or; trained using synthetic images then evaluated on experimentally mixed images (green bars), or; trained using experimentally mixed images then evaluated on experimentally mixed images (blue bars). **(E)** Box plots show the PCC for single-cell features extracted from unmixed images to those extracted from initial single labelling using CellProfiler^31^ ; showing close correspondence between SemaPlex multiplexing and CycIF downstream, at the cell level. **(F)** Median Mean PCC for pixels and features (per marker) compared between initial single labelling (as per A) and unmixed labels, or CycIF relabelling. Correlation between unmixed markers and original labelling is comparable to correlation observed when relabelling and imaging a single molecular target . All error bars/box plots in all panels show lower and upper quartiles for 5-fold cross validated experiments. Significance tested using Welsh’s two-sided t-test. Scale bars = 10μm.

As depicted in **Figure 3A**, for experimental SemaPlex model-training, we have designed a highly efficient, scalable labelling and image acquisition scheme to obtain both individual marker images (deep learning model targets) and mixed marker images (deep learning model inputs) from the same cells. This requires a minimal number of separate cell populations (e.g. wells); equal to the (maximum) number of markers to be unmixed from within each co-labelled channel; here, 3-plex per co-labelled channel, entailing 3 wells. Given this schema, an initial labelling and imaging round acquired each marker individually in its own channel. Next, addition of the two remaining markers in each channel group (excepting the DAPI channel) in each well permitted acquisition of mixed labelling of all three markers per channel in the same image fields (i.e. the same cells). These image sets (individual and mixed markers) were then used to train deep learning models to reconstruct each individual marker from its channel-mixture. In total, nine deep learning models were thus trained (three per channel where mixing was applied). Overall, 49 image fields were acquired per well for mixed markers matched with a single target marker per channel. These 49 image fields were processed into a total of 441 patches (**Methods**). Conducting 5-fold cross validation with an 80:20 train-test splitting, 353 patches were used for training per fold and 88 patches were used for testing per fold. Using mixed marker input images, these models were then applied across all wells to achieve semantic multiplexing of all 10 listed markers in all cells from all wells. Structured in this way, we eliminate the need to generate a separate dataset for the training and application of unmixing models, while also minimising required antibody/label usage. We note that once unmixing models are trained, their further application only requires ‘one-shot’ acquisition of the mixed markers, making this approach readily compatible with high-throughput use.

Purely to verify SemaPlex performance, we also bleached the mixed markers and relabelled each marker individually (1 marker per channel, all 3 wells) using CycIF (over 3 cycles), generating a ground-truth experimentally multiplexed dataset for all markers in all (the same) cells. This facilitated quantitative comparison to the semantically multiplexed dataset at matched single-cell resolution.

### Performance of Semantic Multiplexing on Organelle and Cell Cycle-Associated Markers

Using the SemaPlex dataset detailed above, we compared each unmixed marker to the same marker when initially labelled (rather than the subsequent CycIF labelling). This minimises any confounding effects caused by relabelling differences, as further detailed below. From nine markers imaged in three channels, markers were separated and reconstructed with high visual accuracy (**Figure 3B, C**), achieving Pixel-PCC scores in the range of 0.85 to 0.95 (**Figure 3D**, blue bars), with the exception of ZO-1, which performed relatively poorly (∼0.72) due to very weak labelling signal in the mixed channel. This confirmed our earlier results simulating marker signal imbalances. In further consideration of fundamental, pixel-level performance, we next compared the combined experimental and deep learning strategy of SemaPlex, which promotes efficient training of unmixing models with real experimentally labelled pairs of individual and mixed marker images, to wholly or predominantly synthetic training described for other emerging semantic unmixing implementations^21,23^. Specifically, we compared the Pixel-PCC values for: i) models trained with synthetically mixed markers applied to unmix (unseen) synthetically mixed markers (**Figure 3D**, grey bars) – *very high performance but not real-world*; ii) models trained with synthetically mixed markers applied to unmix (unseen) experimentally mixed (co-labelled) markers (**Figure 3D**, green bars) – *greatly reduced performance indicating mismatch of synthetic training to real-world application*; iii) models trained on and applied to real-world, experimentally mixed marker images (**Figure 3D**, blue bars) – *significant and often pronounced increases in experimentally applied performance, e.g. Pixel-PCC improvements for β-tubulin, p21 and ZO-1 of > 0.3 relative to synthetic training-with-experimental evaluation)*. Thus, while purely synthetic training regimes are extremely convenient and may provide sufficient performance for some applications, results obtained from training using experimentally mixed data can dramatically improve semantic multiplexing results, likely due to fundamental differences in computational *versus* experimental signal-mixing dynamics.

Beyond pixel-level correspondence, we also confirmed that SemaPlex datasets maintained high quantitative fidelity when assessed at single-cell level using gold-standard, quantitative morphometric feature profiling; a typical precursor to downstream biological analyses. To this end, we employed CellPose segmentation^32^ and CellProfiler^31^-based single-cell feature extraction to quantify 133 features per marker, per cell (**Supplementary Table 3**) spanning intensity, radial distribution, and texture characteristics. High correlations quantified between per cell features (termed Feature-PCC) derived from initial single-marker labelling (ground-truth) and matched unmixed markers confirmed recapitulation of quantitative biological information delineated by marker levels and subcellular distributions per cell (**Figure 3E**). Indeed, most markers scored at or above 0.9 for Feature-PCC across intensity, radial distribution, and texture feature-categories at the single-cell scale. To put these values into context, we also measured both Pixel-PCC and Feature-PCC (**Figure 3F**, left and right respectively) between initial marker labelling and subsequent CycIF relabelling of the same marker. Ideally, such repeat labelling would produce identical results, yet factors such as noise in image acquisition and stochastic immunolabelling inevitably cause differences; defining the maximum real-world reproducibility of repeat marker labelling. Accordingly, while relabelling correlations were extremely high across the markers, correlation levels between unmixed markers produced by SemaPlex and initial labelling were comparable. This suggests that inaccuracies associated with SemaPlex approach the margins of error achievable with real-world fluorescence marker labelling and imaging.

### SemaPlex Mimics Biological Resolution of Experimentally Multiplexed Labelling for Cell Cycle Analysis

Again using the dataset defined above, we next sought to visualise single-cell states spanning the cell cycle by performing unsupervised PHATE^33^ manifold embedding based on quantitative marker features derived per cell from cell-matched SemaPlex or CycIF marker image sets (**Figure 4A**). Manifold construction utilised intensity, radial distribution, and texture features measured from cytoplasmic and/or nuclear cell subdomains for each marker (**Supplementary Table 3**), as well as cell morphological features. We observed close correspondence between the ground-truth CycIF manifold and the Semantically Multiplexed manifold, with both elucidating similar biological dynamics. For instance, the colour-coding in both manifolds of Figure 4A indicates three-way relative expression levels of p21, ki67 and PCNA (p21-ki67-PCNA relative expression-level indicated in colour-scale triangle), showing that cells with high p21 expression (associated with suppressed proliferation^34^) cluster at the top of the manifold (marked as milestone 1 in manifold). Cells gradually transition to high PCNA expression, linked to DNA synthesis in interphase^35,36^ (milestone 2). Moving further down, milestones 3 and 4 denote cells high in ki67, whose concentration peaks in late interphase and during mitosis^37^. Per cell expression levels for each of these markers are colour-coded across the same PHATE manifolds in **Figure 4B**, confirming high agreement between CycIF and SemaPlex manifolds in terms of their biological interpretations. Representative SemaPlex cell image examples at each milestone are also visually consistent in marker levels and localisation detail when compared to cell-matched, ground-truth CycIF images, with both image sets revealing expected cell cycle dynamics (**Figure 4C**).

**Figure 4.**
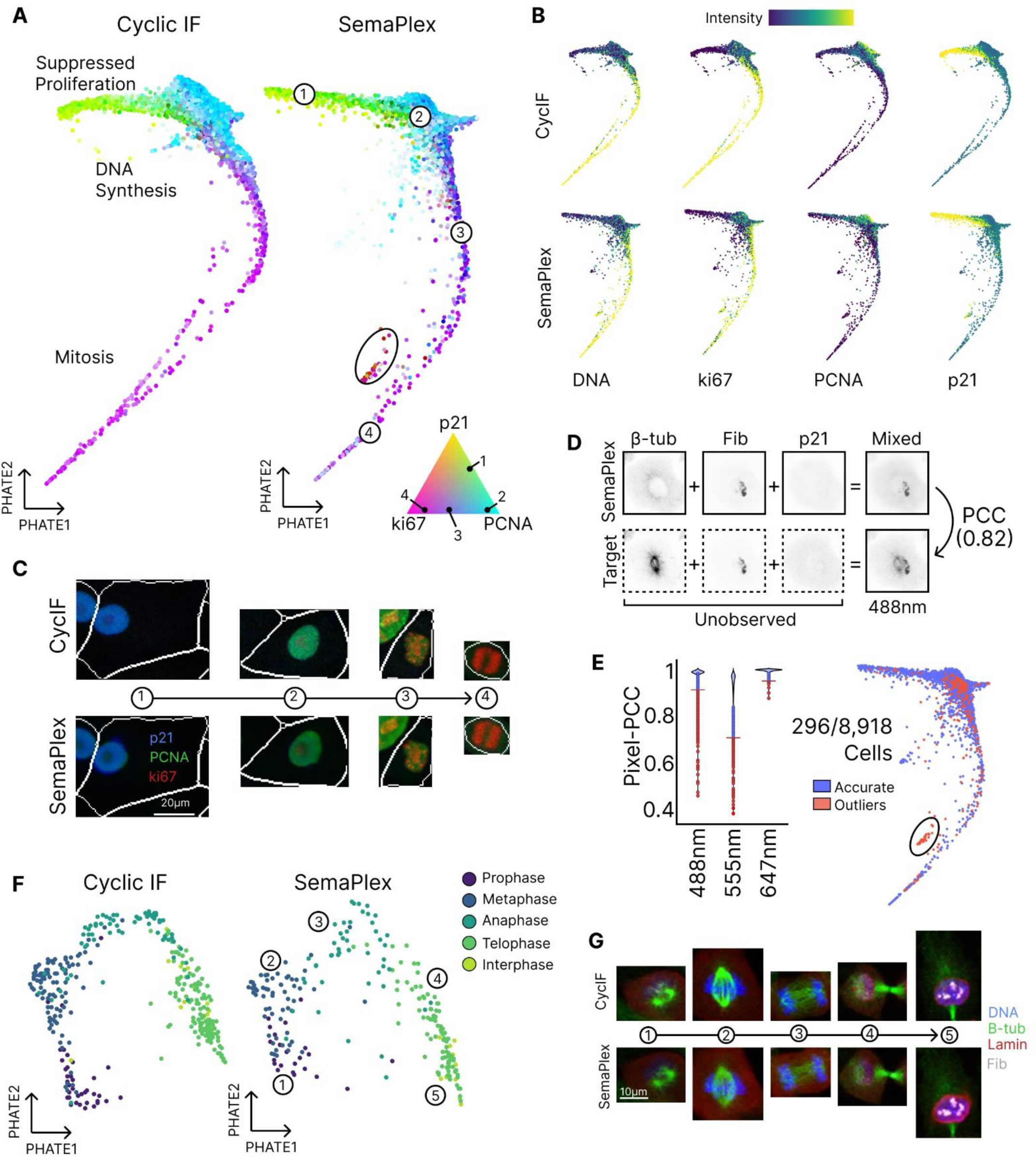
SemaPlex recapitulates cell cycle biology captured by experimental multiplexing. **(A)** CycIF- and SemaPlex-derived single-cell PHATE manifolds constructed using features extracted from 10-marker panel (methods) labelled in A549 cells. Manifolds transition between various expression-mixtures of selected markers, here ki67 (magenta), p21 (yellow) and PCNA (cyan), with marker expression mixtures as per the colour-scale triangle. Numbered milestones mark significant cell state transitions in the cell cycle manifold and correspond to marks on the colour-scale triangle. Examples of cells with aberrant distributions between CycIF and SemaPlex manifolds are circled (and subsequently filtered out as detailed in panel E). **(B)** Selected marker intensities across the PHATE manifolds for key cell cycle markers. **(C)** Representative cells showing key cell cycle markers and morphology matching numbered milestones in A. Composite images contrast-adjusted for display purposes only. Scale bar = 20um. **(D)** For internal detection of erroneous unmixing, unmixed images of markers for a given channel were computationally re - mixed per cell (top row). Pixel-PCC evaluates similarity between the composite image (top row, right) and the original mixed-signal image (bottom row, right). Target images, acquired via CycIF (bottom row, left), are included only to highlight the error detected in unmixing of β-tubulin and are not required for error detection. **(E)** Left: violin plot showing distribution of Pixel-PCC scores per cell when comparing experimentally mixed and computationally re -mixed markers as in D. Thresholds were manually chosen to filter outlier cells with low correspondence and to retain accurate cells. Right: Cells in the SemaPlex manifold highlighted as outliers (red) based on filtering performed for any of the three channels. Circled cells correspond to those outlined in panel A. **(F)** SemaPlex and CycIF manifolds constructed using a subset of cells enriched for mitosis. Cells were manually annotated for mitotic phases, validating unsupervised ordering of cells in both manifolds. **(G)** Representative images along milestones (1-5) from the SemaPlex manifold in F matches the expected mitotic phase progression and corresponding experimental labelling. Images contrast-adjusted for display purposes only. Scale bars = 10μm.

We note that mitotic cells were underrepresented in this realistic application dataset, allowing us to assess SemaPlex robustness to relatively rare cell populations; a common feature of single-cell datasets. To systematically identify cells where semantic multiplexing performed sub-optimally, i.e. to perform ‘error detection’, we recombined (or ‘remixed’) the SemaPlex unmixed markers into a single channel and compared (via Pixel-PCC) that image to the ground-truth experimentally mixed labelling (**Figure 4D**). Marker sets including sub-optimal unmixing achieved low Pixel-PCC values, with Pixel-PCC thresholds applied to exclude outliers or errors with accuracy below this value in any one or more of the fluorescence acquisition channels (**Figure 4E**). Here, exclusion thresholds were chosen by visual inspection, selecting a Pixel-PCC value at which unmixed markers visually diverged from experimental labelling; this is nonetheless adjustable depending on error tolerance. Notably, spuriously unmixed cells identified through this process were indeed inconsistent when compared to either CycIF images or the CycIF-derived PHATE manifold. Demonstrated here in practice, this approach provides a systematic, scalable validation and error-detection method for SemaPlex performance without the need for any additional ground-truth labelling, which may be prohibitive in some applications. Cells showing inconsistencies in their reconstruction can then be parsed out, enhancing the quality and reliability of resulting datasets.

Though a proportion of outliers / errors detected above were, as expected, from the underrepresented mitotic class of cells, we nonetheless found that SemaPlex sufficiently captured mitotic biology despite a generalised training regime (without augmentation or oversampling of mitotic cells). To explore this class further, we selected cells in the lower portion of both CycIF and SemaPlex manifolds; regions that were enriched for mitotic cells (**Supplementary Figure 1**). Previously identified cells that showed evidence of spurious unmixing were excluded from the SemaPlex dataset. Performing PHATE manifold embedding now focused only on mitotic cell populations, we again found that the manifold based on SemaPlex-derived cell features effectively matched the manifold determined from CycIF ground-truth labelling (**Figure 4F**). Annotation of mitotic phases (prophase, metaphase, anaphase, telophase, cytokinesis) showed that both manifolds resolved well-ordered trajectories following expected mitotic progression. We highlight that despite systematic parsing of sub-optimally unmixed cells, which resulted in fewer analysable cells, the mitotic process was still well-sampled by the remaining SemaPlex dataset and closely reflected the underlying biology. This extended to subcellular structural details, with SemaPlex faithfully reconstructing the significant structural changes and sequenced reorganisation associated with mitotic phases, as visually exemplified in **Figure 4G**. For cells in the early stages of mitosis (demarcated by milestone 1 in Figure 4F and G), both SemaPlex and CycIF images detailed the expected breakdown of the nuclear envelope^38^ and nucleoli^39^, along with concentration of microtubule structures at two foci^40^. Throughout the middle mitotic stages (milestones 2, 3), microtubule foci migrate to opposing cell poles, forming mitotic spindles that pull separated chromatids apart. Finally, in later stages (milestones 4, 5), mitotic spindles disassemble and a cleavage furrow formed, denoted by dense microtubule bundles connecting the two daughter cells^41^. Ultimately, nuclear foci reform along with re-assembly of the nuclear envelope, indicating re-entry into interphase^42^.

Taken together, this end-to-end real-world application of SemaPlex demonstrates its ability to efficiently enable multiplexed dataset construction from standard 4-plex fluorescence imaging data, achieving extremely high pixel-level and cell-level fidelity facilitating quantitative single-cell phenotypic analyses of a complex, dynamic, multi-molecular cell biological process.

### Semantic Guidance Can Further Enhance SemaPlex Performance

In many deep learning tasks, the provision of additional ‘context’ has been shown to improve model performance^16,43,44^. Given this, we reasoned that the accurate unmixing of markers achieved by SemaPlex might be further enhanced by providing the deep learning models with extra contextual or semantic cues. We therefore explored a *semantically guided* form of SemaPlex where we augmented mixed marker image inputs with additional channels/markers to improve performance, as depicted in **Figure 1F**. Specifically, we selected ‘challenging’ marker pairs where SemaPlex fidelity was somewhat compromised, such as where unmixing target markers have similar and/or proximal signal patterns / appearance, or where marker signals show very substantial spatial overlap. In the former case, nucleolar marker, fibrillarin, and Golgi complex marker, GM130, are proximal (either within or adjacent to the nucleus, respectively) and show visually similar signal patterns, both being composed of small punctate structures (**Figure 5A**). This proximity and structural similarity results in occasional yet nonetheless systematic unmixing errors close to the nuclear-cytoplasmic interface. We therefore reasoned that semantic guidance from DNA labelling might correct such errors by providing contextual awareness of nuclear location and boundaries. To test this, we applied standard SemaPlex, which we here term *naïve* SemaPlex, by providing only (synthetically) mixed fibrillarin and GM130 images as input for fibrillarin or GM130 unmixing. We then compared performance to a *semantically guided* SemaPlex version which also inputs DNA labelling (DAPI) as a ‘semantic guide’ channel. To evaluate relative unmixing performance of naïve *versus* guided SemaPlex models, we applied Otsu thresholding^45^ to segment fibrillarin (**Figure 5B**) and GM130 (**Supplementary Figure 2A**) signals in real and unmixed images. To generate a single-cell resolved analysis, Cellpose cell segmentation^32^ was then performed based on β-catenin labelling of the same cells (**Supplementary Figure 2B**), before the Jaccard index^46,47^ (measuring segmentation overlap) was calculated per marker, per cell between unmixed marker Otsu segmentation and real (ground-truth) target marker Otsu segmentation. Comparing Jaccard values for naïve *versus* semantically guided SemaPlex at cell population level, we found that guided SemaPlex significantly improved the accuracy of fibrillarin unmixing (**Figure 5C**), particularly by limiting the number of cells with low fidelity (<0.65). This is more clearly shown by comparing per cell Jaccard scores for naïve *versus* guided SemaPlex, where improvements in Jaccard score (values above red diagonals) given guided SemaPlex are particularly common and large for cells with a low Jaccard score after naïve SemaPlex (**Figure 5D**). The detailed nature of these improvements is exemplified in **Figure 5E**, where some punctate GM130 signal was mistaken for fibrillarin with naïve unmixing, whereas the addition of a nuclear marker (DAPI defining DNA) as a semantic guide corrected this error. Similar improvements across various cells are depicted in **Figure 5F**. Repeating the analysis for GM130, we note some similar improvements but these were generally smaller and less common, likely due to the relatively small proportion of (correctable) error compared to the proportion of signal (**Supplementary Figure 2C-E**).

**Figure 5.**
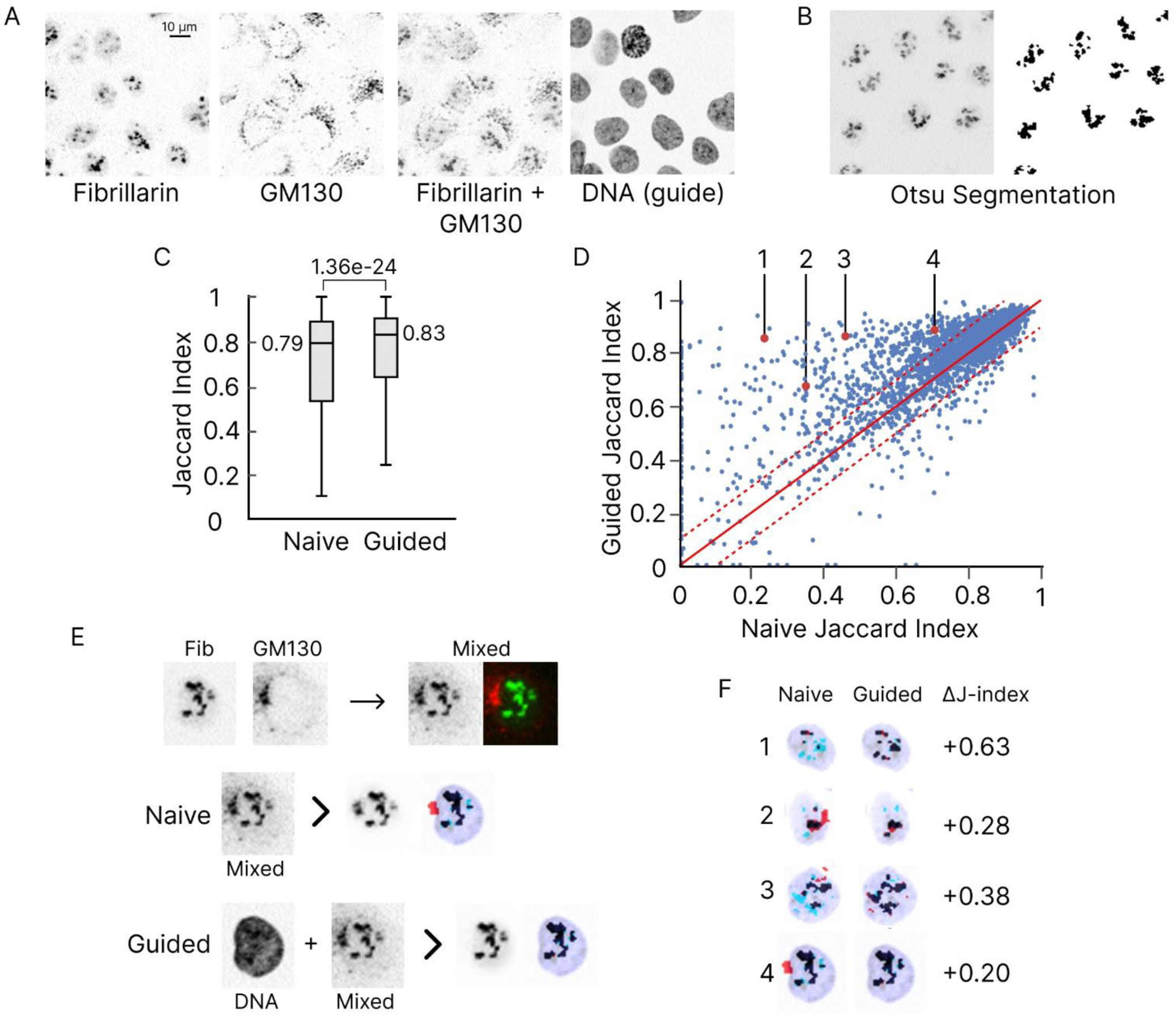
Semantically Guided SemaPlex corrects errors in poorly unmixed cells. (**A**) Example field of fibrillarin, GM130 and their synthetic merge. DAPI-based DNA labelling, which is used as the ‘semantic guide’ channel in this example, is also shown. (**B**) Example segmentation of fibrillarin using Otsu thresholding. (**C**) Population level comparison of naïve *versus* semantically guided SemaPlex performance per cell quantified via the Jaccard index calculated between segmentations of unmixed and real fibrillarin. (**D**) Per cell comparison of naïve *versus* semantically guided Jaccard indexes for fibrillarin segmentation. Solid red line shows where the Jaccard index is equivalent for naïve and guided approaches, and dashed lines show ± 0.1 Jaccard index between the two approaches. (**E**) Example cell with GM130 visually similar to fibrillarin when mixed, as shown in the top row. Middle row shows a naïve fibrillarin reconstruction, with significant over-segmentation (red pixels) where GM130 lies in the mixed channel and outside the nucleus (denoted by the faded blue DNA overlay). Blue pixels denote fibrillarin under-segmentation. The bottom row shows additional semantic guidance provided by the DNA channel, correcting SemaPlex reconstruction of fibrillarin. (**F**) Various cell examples (numbered 1-4 in panel D and depicted as in Panel E) comparing fibrillarin segmentation given naïve *versus* guided SemaPlex. Jaccard index value difference between naïve and semantically guided SemaPlex are shown in the final column. Error bars/box plots in all panels show lower and upper quartiles for 5-fold cross validated experiments. Significance tested using Welsh’s two-sided t-test. Scale bars = 10μm.

An alternative example of semantically guided SemaPlex is also shown for multiplexing of CoxIV (mitochondrial marker) and α-tubulin (microtubule marker); which label cellular structures that are highly overlapping in subcellular space. Here, label-free differential interference contrast (DIC) images were employed as semantic guide channels, and Pixel-PCC of unmixed *versus* ground-truth markers was used to evaluate performance differences between naïve *versus* guided SemaPlex (**Supplementary Figure 2F-I**). While statistically significant improvements in unmixing fidelity were detectable for both markers given semantic guidance, α-tubulin reconstruction was most dramatically improved. This highlights not only the performance improvements accessible with this semantic guidance strategy but also emphasises the latent potential for performance boosts using label-free channels that do not occupy any spectral bandwidth.

## Discussion

Highly multiplexed fluorescence imaging is an immensely powerful technique for spatially resolving the complex, multi-molecular processes that underpin cell biology. To make multiplexed fluorescence imaging far more accessible, we have developed an end-to-end experimental and computational framework, semantic multiplexing (SemaPlex), for efficient amplification of marker detection numbers by several fold. This is accomplished through deep learning-enabled unmixing of multiple molecular markers acquired concurrently within standard fluorescence imaging channels. In this study, we have experimentally shown that SemaPlex achieves not only high visual fidelity, but also faithfully preserves biological content; with both pixel-level and single-cell feature-level correlations to ground-truth approaching the upper limits of reproducibility defined by molecular marker re-labelling in the same cells. Showcased in the contexts of the cell cycle and mitosis biology, SemaPlex resolves marker and morphological dynamics highly consistent with those determined by ground-truth experimentally multiplexed labelling within the same cells (used only for validation independent to SemaPlex labelling). This provides strong real-world evidence for SemaPlex efficacy and its practical application to quantitative single-cell phenotyping of complex biology; demonstrating downstream applied utility that extends upon emerging, largely synthetic evaluations of unmixing focused at the level of pixel similarity rather than cell phenotype fidelity^21,23^.

SemaPlex encompasses a compact and efficient experimental design for the acquisition of cell-matched images of both individual markers and marker-mixtures, enabling generation of real training data representative of actual experimental applications. Importantly, our results indicate that, when applied to real-world datasets, generative unmixing models trained on real experimental data strongly outperform models trained only with synthetic approaches (i.e. simulated marker mixing)^21,23^ (**Figure 3D**). Performance levels from synthetically trained-to-synthetically applied models (only) may thus risk performance-inflation relative to real applications. While this result emphasises the importance of real-world training data for real-world applications, synthetic unmixing experiments are nonetheless valuable, since they allow systematic characterisation of relative performance determinants (e.g. suitable marker combinations, labelling signal requirements, marker numbers) in this new multiplexing paradigm. Taken together, our real-world, experimentally-based analysis of SemaPlex demonstrates an accurate, accessible and scalable workflow for computational multiplexing, whose optimisation can be accelerated with rigorous in-silico testing.

In our analysis of cell cycle-related phenotypes, mitotic cells were a minority and were thus underrepresented in our generalised training. This (intentionally) mirrors imbalanced training that often arises in single-cell data given rare cell subpopulations whose identities are not known *a priori*, or given unique phenotypes produced across large-scale perturbation screening^4,48^. Despite resulting in a relatively higher level of unmixing error in mitotic cells, the inclusion in our SemaPlex approach of an internally validated error detection and tuneable filtration strategy maintains high data quality that reflects the underlying cell biology. Indeed, we find that SemaPlex robustly recapitulates extreme structural variability resulting from sampling of mitosis dynamics; achieving multi-molecular reconstructions that are highly consistent with ground-truth annotations and experimental multiplexing (CycIF) controls. Thus, SemaPlex is effective at handling imbalanced single-cell data distributions even without targeted data augmentation, which is not always possible if cell subpopulations are not defined.

To support our SemaPlex strategy with another innovative layer of performance-enhancement, we assessed the use of ‘semantic guide’ channels, whose inclusion as additional unmixing model-inputs improves model performance by providing added ‘context’; a key factor in model performance across various domains of deep learning^16,43,44^. We exemplify the benefits of this approach by delineating nuclear location using DAPI (DNA) as a semantic guide to correct instances where punctate Golgi signal (GM130) outside of the nucleus is misassigned as punctate nucleolar signal (fibrillarin) inside of the nucleus. Inclusion of the nuclear semantic guide intuitively demonstrates how added spatial context helps the unmixing model to correctly assign the identity of GM130- or fibrillarin-puncta that are otherwise in close proximity and of similar appearance. In an alternative implementation of semantic guidance, we show that label-free imaging modalities (here, DIC) can substantially improve microtubule (β-tubulin) separation from mitochondrial signal (SDHA), despite the absence of explicit human-observable visual cues indicative of this guidance-benefit. Yet this result mirrors recent recognition of the rich latent structural and biological information present in label-free modalities^16,49,50^. While utilisation of semantic guidance markers or channels may rely on availability and experimental context, label-free alternatives are widely compatible with fluorescence imaging, suggesting their generalised suitability as semantic guides for unmixing efforts. Selection of more targeted semantic guides may then additionally benefit from specific domain knowledge and testing to enhance future SemaPlex applications. This provides a significant direction for future development of the SemaPlex approach.

In addition to the impressive quantitative reconstructions of cell cycle and mitotic phenotype heterogeneity demonstrated herein using unsupervised manifold learning (i.e. PHATE), we also demonstrated that semantic multiplexing and individual reconstruction of markers mixed into a single channel can significantly improve deep learning-based classification of subtle (early EMT) cell phenotypes, when compared to classification using the mixed marker images directly (**Figure 2H**). This is of note, since channel ‘overloading’ with multiple mixed markers/dyes is a widely used methodology^13,51^. Notably, despite the same initial content (a single channel containing a mixture of markers), SemaPlex supports even deep learning methods (ResNet18) to more effectively leverage the biological information content of image data and to thereby improve phenotype classification performance. This is likely achieved through the surfacing of latent information present in the images; an effect analogous to feature engineering^16,18,52^ or representation learning^53^. By testing phenotype classification performance across larger numbers of mixed markers (up to 8/channel; at least 2-fold more than other emerging unmixing demonstrations^14,21,23^), we found that while adding more markers improved phenotype classification even without unmixing, performance gains were significantly boosted by applying SemaPlex to separate marker mixtures into their constituent marker signals. More precisely, we found that the benefits of SemaPlex were particularly strong at intermediate numbers of mixed markers (4-6/channel). This appears to reflect competing effects associated with the use of ever higher marker numbers per channel, i.e. increasing total biological information *versus* increasing marker overlap resulting in reduced marker unmixing fidelity. Thus, these results suggest that 4-6 markers per available channel may optimise SemaPlex performance while maintaining labelling efficiency and scalability for even high-throughput phenotypic screening applications. At the same time, since SemaPlex is compatible with complementary spectral and cyclic multiplexing approaches, it can also be viewed as a powerful, customisable strategy through which to dramatically increase phenotyping depth for more targeted, low-throughput applications.

In summary, SemaPlex is an accurate, accessible and scalable method that enables several-fold amplification of molecular marker plexity for fluorescence imaging applications, despite requiring only standard (immuno)fluorescence labelling reagents and hardware. We have shown that semantically multiplexed markers retain biological information comparable to that accessed via experimentally multiplexed imaging, yet SemaPlex carries a significantly reduced experimental burden. SemaPlex thus presents a potent new strategy by which to efficiently capture the rich, spatially resolved multi-molecular data necessary to understand complex cellular processes and states. This makes such data acquisition far more achievable for the researcher community, empowering biologists to conduct more powerful image-based analyses.

## Methods

### Cell culture and imaging

#### Cell culture

HCC38 cells were grown in RPMI 1640 Medium without phenol red (Thermo Fisher Scientific) while A549 cells were maintained in Dulbecco’s Modified Eagle Medium (DMEM, Thermo Fisher Scientific). Media were supplemented with additions as outlined in **Table 1**.

**Table 1:** Media Additions.

| Medium | Supplement | Concentration | Supplier |
| --- | --- | --- | --- |
| RPMI 1640, no phenol red | Fetal bovine serum | 10% (v/v) | Thermo Fisher Scientific |
|  | L-glutamine | 4mM | Merck, Australia |
| DMEM | Fetal bovine serum | 10% (v/v) | Thermo Fisher Scientific |
|  | Penicillin/Streptomycin | 1% (v/v) | Thermo Fisher Scientific |

Cells were cultured at 37°C with 5% CO2 and 21% O2 to 80% confluence before harvesting with 0.05% Trypsin-EDTA (Thermo Fisher Scientific) incubation for 3-5 minutes. 3000 HCC38 cells and 3000 A549 cells were seeded onto (different) 96-well #1.5 glass-bottomed plates (Cellvis) and cultured for 48 hours, generating two separate datasets.

#### EMT Induction, Fixation and Permeabilization

The HCC38 cells were serum-starved for 24 hours and to induce EMT, cells were treated in triplicate with 10 ng/mL of TGF-β1 3 hours, 1 hour, or 20 minutes prior to simultaneous fixation. Both HCC38 cells and A549 cells were fixed using 4% paraformaldehyde (PFA; Electron Microscopy Sciences) for 15 minutes at room temperature (RT), followed by permeabilization with 0.1% Triton X-100 (Thermo Fisher Scientific) for 20 minutes at RT.

#### Immunofluorescent Labelling

A549 cells were used to generate the experimental SemaPlex dataset while HCC38s were used for the EMT-induction dataset. Fixed and permeabilized A549 cells were first pre-treated with a bleaching buffer solution of 20mM NaOH and 3% H2O2 in 1x PBS. Both sets of cells were then blocked for 30 minutes to 1 hour with PBS blocking buffer with 4% w/v bovine serum albumin (BSA; Sigma-Aldrich). Following blocking, cells were incubated with conjugated antibodies for either 12hrs at 4°C, or 2 hours at RT. To label cell nuclei, DAPI was applied for 2hrs at RT alongside antibodies. All antibodies were diluted in 4% BSA and are listed in **Supplementary Table 1.** Cells were washed three times with PBS after fixation and permeabilization, and six times after antibody incubations, and bleaching.

#### Target, Mixed and Cyclic Labelling

For the experimental SemaPlex and CycIF validation datasets, the A549 cells were labelled in 3 distinct rounds: target labelling; mixed labelling; and CycIF labelling. To collect target training images, cells were labelled with DAPI in 405nm, and single markers per channel for 488nm, 555nm and 647nm, with antibody markers in each channel alternating across the three required training wells, as indicated in Figure 3A. Note, the number of wells required for SemaPlex training corresponds to the maximum number of mixed markers to be unmixed per channel; in this case, three. Following image acquisition (detailed below), mixed labelling was performed, involving the addition of antibodies to the appropriate wells to complete mixed channel panels. Specifically, as depicted in Figure 3A, well 1 was labelled with DAPI (405nm), p21 (488nm), Ki67 (555nm), SDHA (647nm), imaged and additionally labelled with Fibrillarin + β-tubulin (488nm), Caveolin-1 + Lamin A/C (555nm), PCNA + ZO-1 (647nm), and reimaged; well 2 was labelled with DAPI (405nm), Fibrillarin (488nm), Caveolin-1 (555nm), PCNA (647nm), imaged and additionally labelled with p21 + β-tubulin (488nm), Ki67 + Lamin A/C (555nm), SDHA + ZO-1 (647nm), and reimaged; well 3 was labelled with DAPI (405nm), β-tubulin (488nm), Lamin A/C (555nm), ZO-1 (647nm), imaged and additionally labelled with p21 + Fibrillarin (488nm), Ki67 + Caveolin-1 (555nm), PCNA + SDHA (647nm), and reimaged. Following the acquisition of mixed images – which completes the requirement for SemaPlex training – CycIF was initiated to generate an additional ground-truth labelling of all markers in all cells; strictly for comparative analysis. Specifically, cells were treated with bleaching buffer solution (detailed previously) and exposed to table lamp illumination for 1 hour at RT. After fluorophore inactivation, CycIF was performed on the sample, involving three cycles of blocking, labelling, imaging and ablation for markers in the 488nm, 555nm and 647nm channels. CycIF was also performed on the EMT-induced HCC38 cells. In each case, CycIF followed the original published methods^7^.

#### Confocal microscopy

All imaging was performed using an AX-R2 confocal microscope (Nikon, Tokyo, Japan) using JOBS automation to ensure reproducible and rapid multi-well, multi-position acquisition. Optimal and readily reproducible focusing was achieved via a combined approach of automated laser-based and image-based focusing using DAPI labelling. For the experimental SemaPlex and CycIF validation datasets, a 40x Plan Apochromat air objective (NA 0.95) was used to acquire images at a spatial resolution of 0.22 μm/pixel. For the EMT-induction dataset, a 20x Plan Apochromat air objective (NA 0.75) was used, with images acquired at spatial resolution of 0.43 μm/pixel. All datasets were imaged at 1x zoom with images of 2048 by 2048 pixels. To ensure consistency, channel-specific optical configurations (OCs) were preserved across target imaging and mixed imaging rounds for SemaPlex. Additionally, the OCs were also maintained across wells, with settings optimal for the brightest marker in each mixed marker channel. For CycIF, OCs were optimised independent of previous SemaPlex OC settings. For quantitative analyses, raw 16-bit .nd2 files were converted to TIFF using custom Python scripts. When required, for visualisation purposes only, image LUTs were adjusted using FIJI software (National Institutes of Health).

#### Image Pre-processing

Image cycles were registered using SIFT features^54^ extracted from DNA labelling which is repeated for every round/cycle. Due to uneven 405nm background illumination in the SemaPlex experiment, we applied field flattening based on BaSiC^55^ for the 405nm channel only. Given insufficient correction, we manually adjusted the correction image to better reflect background illumination curvature, obtaining DAPI images with spatially consistent intensity. Additionally, minor labelling differences between marker mixtures were observed and manually adjusted in subsequent image processing by computationally adding/subtracting a percentage of the initially imaged (individual) marker from the subsequent mixed marker image. Specifically, based on visual inspection, we pixel-wise subtracted 0.5 * PCNA to the mixture in well 1 for the 647nm channel and added 0.3 * β-tubulin to the mixture in well 2 for the 488nm channel. This demonstrates a strategy by which to account for labelling differences across wells and is completely compatible with standard SemaPlex acquisition.

#### SemaPlex Unmixing model design and training

SemaPlex Unmixing was performed using a U-Net^24^ -like architecture with skip connections connecting down convolution blocks to up convolution blocks. 4x4 convolutions, 2x2 stride, and 1x1 padding were used. Leaky ReLU activation with 0.2 negative slope was used for down convolutions and ReLU activation was used for up convolutions. Tanh activation was used after the final up-convolution. Apart from the first and last layer, batch normalisation (ε = 10^-5^, momentum = 0.1) followed activation. Layer size was initialised at 64 filters and doubled until 512 filters. A total of 8 layers were used. 0.5 dropout regularisation was added for 3 of the layers before the deepest layer. L1 loss was used as the objective function, optimised using ADAM^56^ (β_1_ = 0.5, β_2_ = 0.999) for 50 training epochs with a learning rate of 0.0002 before linearly decaying for another 50 epochs. Vertical and/or horizontal flipping augmented training data by a factor of four with image pixel intensities scaled between [-0.5, 0.5]. Training and test sets were independently split, with experiments evaluated using 5-fold cross-validation unless otherwise specified. Model training was conducted using an NVIDIA V100 GPU taking roughly 3 hours per fold.

#### Model evaluation and downstream analysis

Simulated mixed channels were generated by taking the pixel-wise mean across selected channels. Uneven intensities were simulated by first multiplying an attenuating factor for all pixels in a channel before mixing. SemaPlex performance was measured at the image level by calculating the Pearson’s correlation coefficient between corresponding pixels in unmixed and target images (Pixel-PCC).

#### HCC38 Classification following EMT-induction

Image fields were categorised into three classes corresponding to their exposure period following TGF-β1 addition: 20 minutes; 1 hour; or 3 hours. Three wells were imaged for each condition. Images were split into independent training and test sets by holding out images from one well per condition, thereby minimising the potential for information leakage. We trained ResNet18^57^ from scratch for 300 epochs via stochastic gradient descent (learning rate = 0.0001, momentum = 0.9), using a fully connected final layer with 3 output features corresponding to the three classes, activated with the log softmax function.

#### Single-cell analysis using CellProfiler

A549 Cells were segmented with Cellpose^32^ using the default ‘Cells’ model, given DNA (DAPI) and β-tubulin channels. Quantitative feature extraction was then performed using a custom CellProfiler^31^ pipeline. Cells in contact with the image border were excluded, preventing measurement of partial cells. For every cell, 12 morphology features were measured, along with 12 intensity, 69 radial distribution, and 52 texture features for every marker, considering the nuclear (not for radial distribution), cytoplasmic and cell body scales (**Supplementary Table 3**). 8,918 A549 cells were analysed from a single biological replicate across three wells with 350 non-mitotic multi-nucleated cells excluded from analysis. Nuclear features for mitotic cells with multiple ‘nuclei’ were summed for integrated features or averaged otherwise. All features were standard-scaled per well. Unsupervised embeddings were created via PHATE^33^ with KNN=20 for manifolds containing all cells and KNN=12 for manifolds containing mitotic cells. Embeddings utilised morphology features, as well as intensity, radial distribution, and texture features from all 10 markers labelled at the nuclear scale, and at the cytoplasmic scale for β-tubulin, ZO1, SDHA, and CAV-1.

Internally validated SemaPlex error-detection was for performed per cell by comparing experimentally mixed labelling (used as input for unmixing models) to unmixed labels remixed back into a single channel. For good unmixing, both images should be broadly equivalent, measured by Pixel-PCC. Since relative intensities for remixing were unknown *a priori*, we optimised the Pixel-PCC between experimentally mixed image and a linear combination of unmixed channels. To filter out poorly unmixed cells, we set a threshold 0.9, 0.75, 0.95 Pixel-PCC for 488nm, 555nm, 647nm channels, respectively. These values were manual chosen by visually inspecting SemaPlex labelling for a range of Pixel-PCCs. Cells failing to score above the set thresholds in any channel were considered erroneous, resulting in the exclusion of 296 cells after SemaPlex.

Annotation of mitotic cells were performed manually using DNA, β-tubulin, and Lamin A/C. Cells were classified as prophase, metaphase, anaphase, telophase, or non-mitotic. Cells that could not be confidently classified were excluded from analysis of the mitotic population. Quantitative selection of mitotic-enriched cell populations was manually performed as depicted in **Supplementary Figure 1**.

#### Guided SemaPlex application and analysis

DU145 cells labelling and imaged using the 4i method^6^ from Gunawan *et al.*^16^ were used in this section. Mixed channels were generated computationally as described above. The SemaPlex deep learning model architecture and training process were as described above. Segmentation of fibrillarin and GM130 was performed using Otsu thresholding^45^ leveraging the *skimage* package^58^. Similarly, the Jaccard index^46^ for comparing segmentation profiles was calculated using the *skimage* package. Local Pixel-PCC scores are shown for α-tubulin and CoxIV, based on calculating PCC inside a 3x3 pixel sliding kernel, with images undergoing reflective padding to ensure complete border coverage of local PCC estimates.

#### Statistics & Reproducibility

Computation experiments subject to statistical testing were repeated using 5-fold cross-validation with significance tests conducted using Welsh’s two-sided t-test (unequal variance assumptions) as noted in relevant figure legends. P-value < 0.05 was considered as statistically significant. No data was excluded unless specifically noted.

## Data Availability

Images and CellProfiler outputs generated in this study have been deposited at https://zenodo.org/records/17118601.

## Code Availability

Code used to train and apply models is made available at https://github.com/CancerSystemsMicroscopyLab/SemaPlex. Sample scripts used to generate results in this manuscript are also provided at https://zenodo.org/records/17118601.

## Acknowledgements

The authors extend special thanks to Dr Sian Culley for valuable feedback on the manuscript. I.G. and F.V.K. are supported by Australian Government Research Training Program (RTP) Scholarships. M.D. is supported by the UNSW University Postgraduate Award. F.V.K. receives a Top-Up award from SPHERE Cancer CAG and Cancer Institute NSW. D.P.N. is supported by a Tour de Cure Pioneering Grant (RSP-573-2024). J.G.L. is supported by a University of New South Wales Scientia Research Fellowship, a Ramaciotti Biomedical Research Award, an ARC Development Project grant (DP170103599), NHMRC Ideas Grants (GNT1184009, GNT2012848, GNT2028506), and a Tour de Cure Pioneering Grant (RSP-547-FY2023). This research was undertaken with the assistance of resources and services from the National Computational Infrastructure (NCI), which is supported by the Australian Government.

## Author contributions

I.G., M.D., J.G.L. conceived the paper and project design. F.V.K and D.P.N contributed to early testing and validation. Unmixing and all downstream analyses conducted by I.G, with assistance from M.D. and Y.H. Preparation, multiplexed labelling and imaging of HCC38 cells conducted by D.P.N. Preparation, labelling, and imaging of multiplexed and 4-plexed A549 cells conducted by M.D. Design and writing of the manuscript was led by I.G, M.D. and J.G.L. Significant conceptual development and editing by E.M.

## References

1. Karacosta, L. G. From imaging a single cell to implementing precision medicine: an exciting new era. Emerg. Top. Life Sci. 5, 837–847 (2021).

2. Lock, J. G. & Strömblad, S. Systems microscopy: an emerging strategy for the life sciences. Exp. Cell Res. 316, 1438–1444 (2010).

3. Pratapa, A., Doron, M. & Caicedo, J. C. Image-based cell phenotyping with deep learning. Curr. Opin. Chem. Biol. 65, 9–17 (2021).

4. Bryce, N. S. et al. High-content imaging of unbiased chemical perturbations reveals that the phenotypic plasticity of the actin cytoskeleton is constrained. Cell Syst. 9, 496–507 (2019).

5. Lichtman, J. W. & Conchello, J.-A. Fluorescence microscopy. Nat. Methods 2, 910–919 (2005).

6. Kramer, B. A., Del Castillo, J. S., Pelkmans, L. & Gut, G. Iterative indirect immunofluorescence imaging (4i) on adherent cells and tissue sections. Bio-Protoc. 13, e4712 (2023).

7. Lin, J., Fallahi-Sichani, M., Chen, J. & Sorger, P. K. Cyclic immunofluorescence (CycIF), a highly multiplexed method for single-cell imaging. Curr. Protoc. Chem. Biol. 8, 251–264 (2016).

8. Gut, G., Herrmann, M. D. & Pelkmans, L. Multiplexed protein maps link subcellular organization to cellular states. Science 361, eaar7042 (2018).

9. Kramer, B. A., Sarabia del Castillo, J. & Pelkmans, L. Multimodal perception links cellular state to decision-making in single cells. Science 377, 642–648 (2022).

10. Lin, J.-R. et al. Multiplexed 3D atlas of state transitions and immune interaction in colorectal cancer. Cell 186, 363–381 (2023).

11. Seo, J. et al. PICASSO allows ultra-multiplexed fluorescence imaging of spatially overlapping proteins without reference spectra measurements. Nat. Commun. 13, 2475 (2022).

12. Gustafsdottir, S. M. et al. Multiplex cytological profiling assay to measure diverse cellular states. PloS One 8, e80999 (2013).

13. Bray, M.-A. et al. Cell Painting, a high-content image-based assay for morphological profiling using multiplexed fluorescent dyes. Nat. Protoc. 11, 1757–1774 (2016).

14. von Coburg, E. et al. Cell Painting PLUS: expanding the multiplexing capacity of Cell Painting-based phenotypic profiling using iterative staining-elution cycles. Nat. Commun. 16, 3857 (2025).

15. Ounkomol, C., Seshamani, S., Maleckar, M. M., Collman, F. & Johnson, G. R. Label-free prediction of three-dimensional fluorescence images from transmitted-light microscopy. Nat. Methods 15, 917–920 (2018).

16. Gunawan, I. et al. Extensible Immunofluorescence (ExIF) accessibly generates high-plexity datasets by integrating standard 4-plex imaging data. Nat. Commun. 16, 1–15 (2025).

17. Sun, G., Liu, S., Shi, C., Liu, X. & Guo, Q. 3DCNAS: A universal method for predicting the location of fluorescent organelles in living cells in three-dimensional space. Exp. Cell Res. 433, 113807 (2023).

18. Lefebvre, A. E. et al. Nellie: automated organelle segmentation, tracking and hierarchical feature extraction in 2D/3D live-cell microscopy. Nat. Methods 1–13 (2025).

19. Zhanghao, K. et al. Fast segmentation and multiplexing imaging of organelles in live cells. Nat. Commun. 16, 2769 (2025).

20. Ben-Uri, R. et al. High-dimensional imaging using combinatorial channel multiplexing and deep learning. Nat. Biotechnol. (2025) doi:10.1038/s41587-025-02585-0.

21. Ashesh, A. et al. Micro𝕊plit: Semantic Unmixing of Fluorescent Microscopy Data. bioRxiv 2025–02 (2025).

22. Laissue, P. P., Alghamdi, R. A., Tomancak, P., Reynaud, E. G. & Shroff, H. Assessing phototoxicity in live fluorescence imaging. Nat. Methods 14, 657–661 (2017).

23. Chen, M., et al. Interpretable deep learning illuminates multiple structures fluorescence imaging: a path toward trustworthy artificial intelligence in microscopy. *ArXiv Prepr. ArXiv250105490* (2025).

24. Ronneberger, O., Fischer, P. & Brox, T. U-net: Convolutional networks for biomedical image segmentation. in 234–241 (Springer, 2015).

25. Pearson, K. LIII. On lines and planes of closest fit to systems of points in space. Lond. Edinb. Dublin Philos. Mag. J. Sci. 2, 559–572 (1901).

26. Gunawan, I. et al. Image quality metrics fail to accurately represent biological information in fluorescence microscopy. bioRxiv 2025–08 (2025).

27. Xu, J., Lamouille, S. & Derynck, R. TGF-β-induced epithelial to mesenchymal transition. Cell Res. 19, 156–172 (2009).

28. Zavadil, J. & Böttinger, E. P. TGF-β and epithelial-to-mesenchymal transitions. Oncogene 24, 5764– 5774 (2005).

29. Stallaert, W. et al. The structure of the human cell cycle. Cell Syst. 13, 230–240.e3 (2022).

30. Carlton, J. G., Jones, H. & Eggert, U. S. Membrane and organelle dynamics during cell division. Nat. Rev. Mol. Cell Biol. 21, 151–166 (2020).

31. Carpenter, A. E. et al. CellProfiler: image analysis software for identifying and quantifying cell phenotypes. Genome Biol. 7, R100 (2006).

32. Stringer, C., Wang, T., Michaelos, M. & Pachitariu, M. Cellpose: a generalist algorithm for cellular segmentation. Nat. Methods 18, 100–106 (2021).

33. Moon, K. R. et al. Visualizing structure and transitions in high-dimensional biological data. Nat. Biotechnol. 37, 1482–1492 (2019).

34. Karimian, A., Ahmadi, Y. & Yousefi, B. Multiple functions of p21 in cell cycle, apoptosis and transcriptional regulation after DNA damage. DNA Repair 42, 63–71 (2016).

35. Moldovan, G.-L., Pfander, B. & Jentsch, S. PCNA, the maestro of the replication fork. Cell 129, 665– 679 (2007).

36. Schönenberger, F., Deutzmann, A., Ferrando-May, E. & Merhof, D. Discrimination of cell cycle phases in PCNA-immunolabeled cells. BMC Bioinformatics 16, 180 (2015).

37. Sobecki, M. et al. Cell-Cycle Regulation Accounts for Variability in Ki-67 Expression Levels. Cancer Res. 77, 2722–2734 (2017).

38. Vivante, A., Shoval, I. & Garini, Y. The dynamics of lamin a during the cell cycle. Front. Mol. Biosci. 8, 705595 (2021).

39. Soldani, C., Bottone, M. G., Pellicciari, C. & Scovassi, A. I. Nucleolus disassembly in mitosis and apoptosis: dynamic redistribution of phosphorylated-c-Myc, fibrillarin and Ki-67. Eur. J. Histochem. 50, 273–280 (2006).

40. Scholey, J. M., Rogers, G. C. & Sharp, D. J. Mitosis, microtubules, and the matrix. J. Cell Biol. 154, 261 (2001).

41. Carlton, J. G., Jones, H. & Eggert, U. S. Membrane and organelle dynamics during cell division. Nat. Rev. Mol. Cell Biol. 21, 151–166 (2020).

42. Chaudhary, N. & Courvalin, J.-C. Stepwise reassembly of the nuclear envelope at the end of mitosis. J. Cell Biol. 122, 295–306 (1993).

43. Roy, S. K. et al. Multimodal fusion transformer for remote sensing image classification. IEEE Trans. Geosci. Remote Sens. 61, 1–20 (2023).

44. Liu, W., Zheng, W.-L. & Lu, B.-L. Emotion recognition using multimodal deep learning. in 521–529 (Springer, 2016).

45. Yousefi, J. Image binarization using Otsu thresholding algorithm. Ont. Can. Univ. Guelph 10, (2011).

46. Ge, F., Wang, S. & Liu, T. New benchmark for image segmentation evaluation. J. Electron. Imaging 16, 033011–033011 (2007).

47. Reinke, A. et al. Understanding metric-related pitfalls in image analysis validation. Nat. Methods 21, 182–194 (2024).

48. Ramezani, M. et al. A genome-wide atlas of human cell morphology. Nat. Methods 22, 621–633 (2025).

49. Rappez, L., Rakhlin, A., Rigopoulos, A., Nikolenko, S. & Alexandrov, T. DeepCycle reconstructs a cyclic cell cycle trajectory from unsegmented cell images using convolutional neural networks. Mol. Syst. Biol. 16, e9474 (2020).

50. Christiansen, E. M. et al. In silico labeling: predicting fluorescent labels in unlabeled images. Cell 173, 792–803 (2018).

51. Sheard, T. M. et al. Differential labelling of human sub-cellular compartments with fluorescent dye esters and expansion microscopy. Nanoscale 15, 18489–18499 (2023).

52. Zheng, A. & Casari, A. Feature Engineering for Machine Learning: Principles and Techniques for Data Scientists. (O’Reilly Media, Inc., 2018).

53. Gunawan, I., Vafaee, F., Meijering, E. & Lock, J. G. An introduction to representation learning for single-cell data analysis. *Cell Rep*. Methods 3, (2023).

54. Lowe, G. Sift-the scale invariant feature transform. Int J 2, 2 (2004).

55. Peng, T. et al. A BaSiC tool for background and shading correction of optical microscopy images. Nat. Commun. 8, 14836 (2017).

56. Kingma, D. P. & Ba, J. Adam: A method for stochastic optimization. *ArXiv Prepr. ArXiv14126980* (2014).

57. He, K., Zhang, X., Ren, S. & Sun, J. Deep residual learning for image recognition. in 770–778 (2016).

58. Van der Walt, S. et al. scikit-image: image processing in Python. PeerJ 2, e453–e453 (2014).

