## Supplementary Data for "Generative semantic multiplexing (SemaPlex) for accessible and scalable multiplexed fluorescence imaging"

Supplementary Table 1. List of antibodies and cell-labelling reagents used.

| | | Conjugate | Supplier | Catalogue number |
|---|---|---|---|---|
| **Conjugated antibodies** | Conc. | | | |
| Pan-cytokeratin | 1:500 | AF488 | Cell Signaling Technology | 8956S |
| Vimentin | 1:200 | AF647 | Biolegend | 677807 |
| psmad2/psmad 3 | 1:200 | AF647 | BD Biosciences | 072-670 |
| CD44 | 1:500 | PE | Cell Signaling Technology | 8724S |
| α-tubulin | 1:500 | AF488 | Thermo-Fisher Scientific | 322588 |
| β-catenin (1) | 1:200 | AF555 | Thermo-Fisher Scientific | 71-2700 A20187 |
| Fibrillarin | 1:500 | AF555 | ABCAM | AB203410 |
| β-tubulin | 1:500 | AF488 | ABCAM | EPR16774 |
| p21 Waf1/Cip1 | 1:200 | AF488 | Cell Signaling Technology | 12D1 |
| Fibrillarin | 1:8000 | AF488 | ABCAM | EPR10823b |
| Lamin A/C | 1:4000 | PE | ABCAM | ab210433 |
| Caveolin 1 | 1:500 | PE | ABCAM | ab212007 |
| Ki67 | 1:500 | AF555 | Cell Signaling Technology | 20701S |
| SDHA | 1:500 | AF647 | ABCAM | AB168536 |
| PCNA | 1:200 | AF647 | Cell Signaling Technology | 82968S |
| ZO-1 | 1:200 | AF647 | Cell Signaling Technology | 98225S |
| **Fluorescent Labels** | | | | |
| 4', 6-diamidino-2-phenylindole (DAPI) | 1:2000 | | Sigma-Aldrich Pty Ltd | D9532 |

(1) Primary antibody was conjugated in-house.

Supplementary Table 2. Marker combinations used in synthetic unmixing evaluation

| # Markers | Markers | Classification |
|---|---|---|
| 2 | Cytokeratin, Vim | Yes |
| | β-catenin, pSMAD2/3 | No |
| | α -tubulin, β-catenin | Yes |
| | β-catenin, Cytokeratin | No |
| | β-catenin, CD44 | No |
| | α -tubulin, Cytokeratin | No |
| | CD44, Vim | Yes |
| | β-catenin, Fibrillarin | No |
| | α -tubulin, Fibrillarin | No |
| | pSMAD2/3, Vim | Yes |
| | DAPI, Fibrillarin | Yes |
| 3 | Fibrillarin, pSMAD2/3, Vim | No |
| | β-catenin, Cytokeratin, Fibrillarin | Yes |
| | β-catenin, DAPI, Fibrillarin | Yes |
| | β-catenin, Cytokeratin, Vim | No |
| | α -tubulin, DAPI, Fibrillarin | No |
| | α -tubulin, CD44, DAPI | Yes |
| | Cytokeratin, pSMAD2/3, Vim | No |
| | CD44, pSMAD2/3, Vim | Yes |
| 4 | α -tubulin, CD44, DAPI, Vim | No |
| | CD44, Cytokeratin, pSMAD2/3, Vim | No |
| | β-catenin, Cytokeratin, Fibrillarin, Vim | No |
| | β-catenin, Fibrillarin, pSMAD2/3, Vim | No |
| | β-catenin, Cytokeratin, Fibrillarin, pSMAD2/3 | Yes |
| | α -tubulin, β-catenin, DAPI, Fibrillarin | Yes |
| | β-catenin, CD44, pSMAD2/3, Vim | Yes |
| 5 | β-catenin, CD44, Fibrillarin, pSMAD2/3, Vim | Yes |
| | β-catenin, Cytokeratin, DAPI, Fibrillarin, Vim | No |
| | α -tubulin, CD44, DAPI, Fibrillarin, Vim | No |
| | β-catenin, Cytokeratin, Fibrillarin, pSMAD2/3, Vim | Yes |
| | α -tubulin, β-catenin, DAPI, Fibrillarin, Vim | Yes |
| 6 | α -tubulin, CD44, Cytokeratin, DAPI, Fibrillarin, Vim | No |
| | β-catenin, CD44, Cytokeratin, Fibrillarin, pSMAD2/3, Vim | Yes |
| | β-catenin, Cytokeratin, DAPI, Fibrillarin, pSMAD2/3, Vim | Yes |
| | α -tubulin, β-catenin, CD44, DAPI, Fibrillarin, Vim | Yes |
| 7 | α -tubulin, β-catenin, Cytokeratin, DAPI, Fibrillarin, pSMAD2/3, Vim | Yes |
| | α -tubulin, β-catenin, CD44, Cytokeratin, DAPI, Fibrillarin, Vim | No |
| | α -tubulin, β-catenin, CD44, DAPI, Fibrillarin, pSMAD2/3, Vim | Yes |
| 8 | α -tubulin, β-catenin, CD44, Cytokeratin, DAPI, Fibrillarin, pSMAD2/3, Vim | Yes |

Supplementary Table 3. List of per cell features measured via CellProfiler.

| **Morphology Features** | **Radial Distribution Features** | **Texture Features** |
|---|---|---|
| 1. AreaShape_Area | 1. RadialDistribution_FracAtD_1of3 | 1. Texture_AngularSecondMoment_3_00_256 |
| 2. AreaShape_Compactness | 2. RadialDistribution_FracAtD_2of3 | 2. Texture_AngularSecondMoment_3_01_256 |
| 3. AreaShape_Eccentricity | 3. RadialDistribution_FracAtD_3of3 | 3. Texture_AngularSecondMoment_3_02_256 |
| 4. AreaShape_Extent | 4. RadialDistribution_MeanFrac_1of3 | 4. Texture_AngularSecondMoment_3_03_256 |
| 5. AreaShape_FormFactor | 5. RadialDistribution_MeanFrac_2of3 | 5. Texture_Contrast_3_00_256 |
| 6. AreaShape_MeanRadius | 6. RadialDistribution_MeanFrac_3of3 | 6. Texture_Contrast_3_01_256 |
| | 7. RadialDistribution_RadialCV_1of3 | 7. Texture_Contrast_3_02_256 |
| | 8. RadialDistribution_RadialCV_2of3 | 8. Texture_Contrast_3_03_256 |
| **Intensity Features** | 9. RadialDistribution_RadialCV_3of3 | 9. Texture_Correlation_3_00_256 |
| | 10. RadialDistribution_ZernikeMagnitude_0_0 | 10. Texture_Correlation_3_01_256 |
| 1. Intensity_IntegratedIntensityEdge | 11. RadialDistribution_ZernikeMagnitude_1_1 | 11. Texture_Correlation_3_02_256 |
| 2. Intensity_IntegratedIntensity | 12. RadialDistribution_ZernikeMagnitude_2_0 | 12. Texture_Correlation_3_03_256 |
| 3. Intensity_LowerQuartileIntensity | 13. RadialDistribution_ZernikeMagnitude_2_2 | 13. Texture_DifferenceEntropy_3_00_256 |
| 4. Intensity_MADIntensity | 14. RadialDistribution_ZernikeMagnitude_3_1 | 14. Texture_DifferenceEntropy_3_01_256 |
| 5. Intensity_MassDisplacement | 15. RadialDistribution_ZernikeMagnitude_3_3 | 15. Texture_DifferenceEntropy_3_02_256 |
| 6. Intensity_MaxIntensityEdge | 16. RadialDistribution_ZernikeMagnitude_4_0 | 16. Texture_DifferenceEntropy_3_03_256 |
| 7. Intensity_MaxIntensity | 17. RadialDistribution_ZernikeMagnitude_4_2 | 17. Texture_DifferenceVariance_3_00_256 |
| 8. Intensity_MeanIntensityEdge | 18. RadialDistribution_ZernikeMagnitude_4_4 | 18. Texture_DifferenceVariance_3_01_256 |
| 9. Intensity_MeanIntensity | 19. RadialDistribution_ZernikeMagnitude_5_1 | 19. Texture_DifferenceVariance_3_02_256 |
| 10. Intensity_MedianIntensity | 20. RadialDistribution_ZernikeMagnitude_5_3 | 20. Texture_DifferenceVariance_3_03_256 |
| 11. Intensity_MinIntensityEdge | 21. RadialDistribution_ZernikeMagnitude_5_5 | 21. Texture_Entropy_3_00_256 |
| 12. Intensity_MinIntensity | 22. RadialDistribution_ZernikeMagnitude_6_0 | 22. Texture_Entropy_3_01_256 |
| 13. Intensity_StdIntensityEdge | 23. RadialDistribution_ZernikeMagnitude_6_2 | 23. Texture_Entropy_3_02_256 |
| 14. Intensity_StdIntensity | 24. RadialDistribution_ZernikeMagnitude_6_4 | 24. Texture_Entropy_3_03_256 |
| 15. Intensity_UpperQuartileIntensity | 25. RadialDistribution_ZernikeMagnitude_6_6 | 25. Texture_InfoMeas1_3_00_256 |
| | 26. RadialDistribution_ZernikeMagnitude_7_1 | 26. Texture_InfoMeas1_3_01_256 |
| | 27. RadialDistribution_ZernikeMagnitude_7_3 | 27. Texture_InfoMeas1_3_02_256 |
| | 28. RadialDistribution_ZernikeMagnitude_7_5 | 28. Texture_InfoMeas1_3_03_256 |
| | 29. RadialDistribution_ZernikeMagnitude_7_7 | 29. Texture_InfoMeas2_3_00_256 |
| | 30. RadialDistribution_ZernikeMagnitude_8_0 | 30. Texture_InfoMeas2_3_01_256 |
| | 31. RadialDistribution_ZernikeMagnitude_8_2 | 31. Texture_InfoMeas2_3_02_256 |
| | 32. RadialDistribution_ZernikeMagnitude_8_4 | 32. Texture_InfoMeas2_3_03_256 |
| | 33. RadialDistribution_ZernikeMagnitude_8_6 | 33. Texture_InverseDifferenceMoment_3_00_256 |
| | 34. RadialDistribution_ZernikeMagnitude_8_8 | 34. Texture_InverseDifferenceMoment_3_01_256 |
| | 35. RadialDistribution_ZernikeMagnitude_9_1 | 35. Texture_InverseDifferenceMoment_3_02_256 |
| | 36. RadialDistribution_ZernikeMagnitude_9_3 | 36. Texture_InverseDifferenceMoment_3_03_256 |
| | 37. RadialDistribution_ZernikeMagnitude_9_5 | 37. Texture_SumAverage_3_00_256 |
| | 38. RadialDistribution_ZernikeMagnitude_9_7 | 38. Texture_SumAverage_3_01_256 |
| | 39. RadialDistribution_ZernikeMagnitude_9_9 | 39. Texture_SumAverage_3_02_256 |
| | 40. RadialDistribution_ZernikePhase_0_0 | 40. Texture_SumAverage_3_03_256 |
| | 41. RadialDistribution_ZernikePhase_1_1 | 41. Texture_SumEntropy_3_00_256 |
| | 42. RadialDistribution_ZernikePhase_2_0 | 42. Texture_SumEntropy_3_01_256 |
| | 43. RadialDistribution_ZernikePhase_2_2 | 43. Texture_SumEntropy_3_02_256 |
| | 44. RadialDistribution_ZernikePhase_3_1 | 44. Texture_SumEntropy_3_03_256 |
| | 45. RadialDistribution_ZernikePhase_3_3 | 45. Texture_SumVariance_3_00_256 |
| | 46. RadialDistribution_ZernikePhase_4_0 | 46. Texture_SumVariance_3_01_256 |
| | 47. RadialDistribution_ZernikePhase_4_2 | 47. Texture_SumVariance_3_02_256 |
| | 48. RadialDistribution_ZernikePhase_4_4 | 48. Texture_SumVariance_3_03_256 |
| | 49. RadialDistribution_ZernikePhase_5_1 | 49. Texture_Variance_3_00_256 |
| | 50. RadialDistribution_ZernikePhase_5_3 | 50. Texture_Variance_3_01_256 |
| | 51. RadialDistribution_ZernikePhase_5_5 | 51. Texture_Variance_3_02_256 |
| | 52. RadialDistribution_ZernikePhase_6_0 | 52. Texture_Variance_3_03_256 |
| | 53. RadialDistribution_ZernikePhase_6_2 | |
| | 54. RadialDistribution_ZernikePhase_6_4 | |
| | 55. RadialDistribution_ZernikePhase_6_6 | |
| | 56. RadialDistribution_ZernikePhase_7_1 | |
| | 57. RadialDistribution_ZernikePhase_7_3 | |
| | 58. RadialDistribution_ZernikePhase_7_5 | |
| | 59. RadialDistribution_ZernikePhase_7_7 | |
| | 60. RadialDistribution_ZernikePhase_8_0 | |
| | 61. RadialDistribution_ZernikePhase_8_2 | |
| | 62. RadialDistribution_ZernikePhase_8_4 | |
| | 63. RadialDistribution_ZernikePhase_8_6 | |
| | 64. RadialDistribution_ZernikePhase_8_8 | |
| | 65. RadialDistribution_ZernikePhase_9_1 | |
| | 66. RadialDistribution_ZernikePhase_9_3 | |
| | 67. RadialDistribution_ZernikePhase_9_5 | |
| | 68. RadialDistribution_ZernikePhase_9_7 | |
| | 69. RadialDistribution_ZernikePhase_9_9 | |

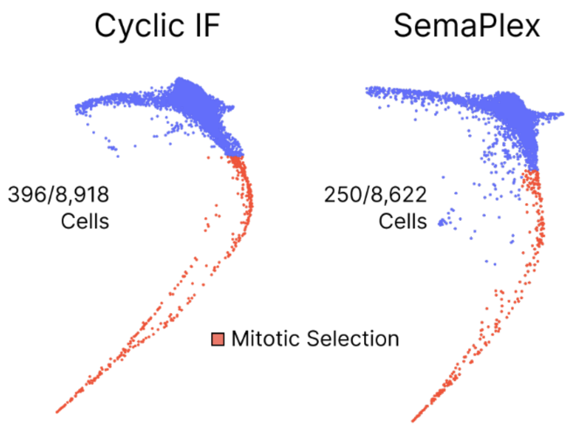


**Supplementary Figure 1.** Selection of mitotic (red) *versus* non-mitotic (blue) cells in PHATE manifolds based either on real multiplexed data (left; CyclicIF) or unmixed data after removal of poorly unmixed cells (right; SemaPlex). Mitotic cell numbers (396 or 250, respectively) out of all cells (8,918 or 8,622, respectively) are indicated.

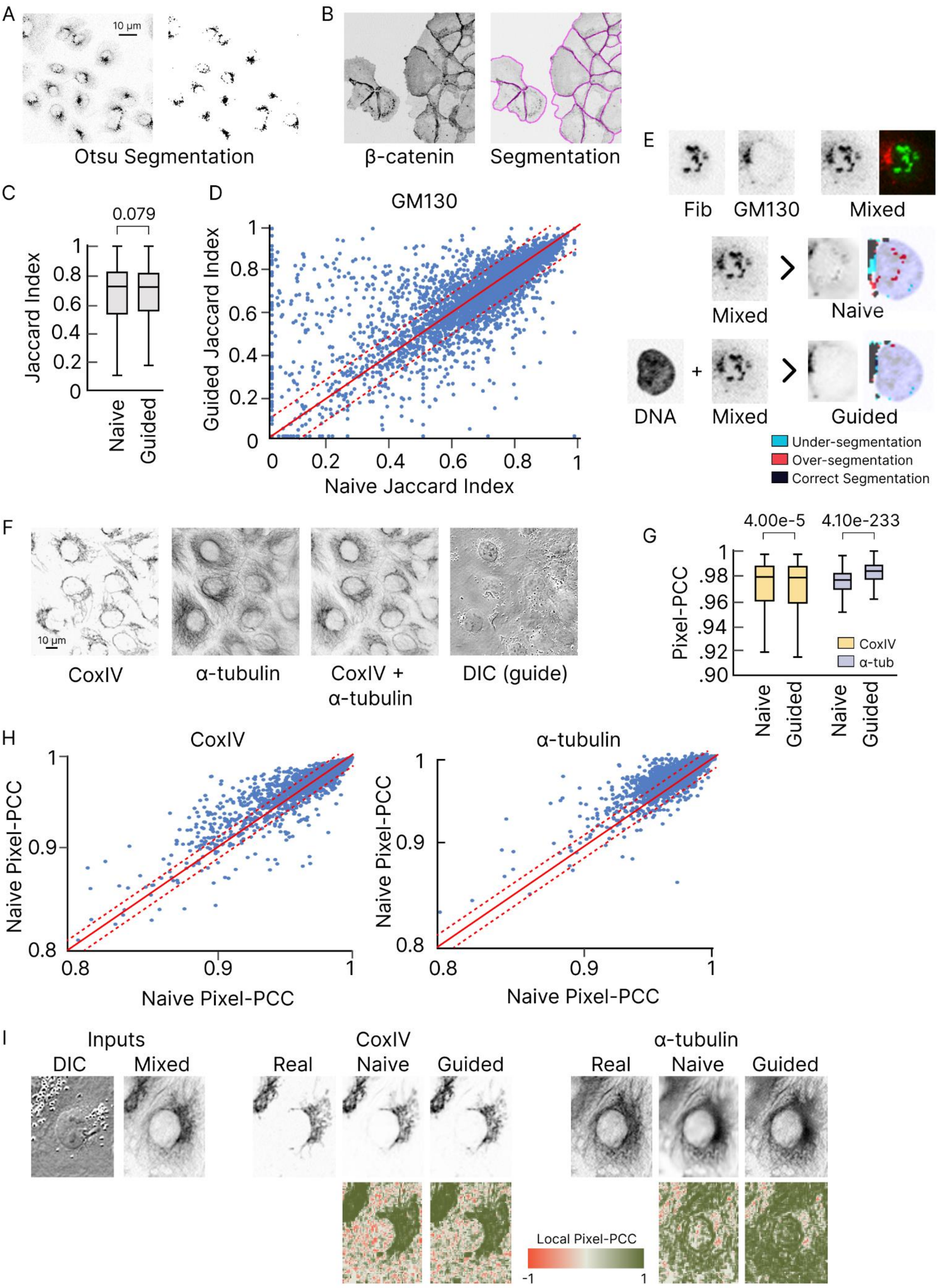
A
10 µm
Otsu Segmentation
B
β-catenin
Segmentation
C
0.079
Jaccard Index
Naive
Guided
D
GM130
Guided Jaccard Index
Naive Jaccard Index
E
Fib
GM130
Mixed
Mixed
Naive
DNA
Mixed
Guided
Under-segmentation
Over-segmentation
Correct Segmentation
F
10 µm
CoxIV
α-tubulin
CoxIV + α-tubulin
DIC (guide)
G
4.00e-5
4.10e-233
Pixel-PCC
CoxIV
α-tub
Naive
Guided
Naive
Guided
H
CoxIV
α-tubulin
Naive Pixel-PCC
Naive Pixel-PCC
Naive Pixel-PCC
Naive Pixel-PCC
I
Inputs
DIC
Mixed
CoxIV
Real
Naive
Guided
α-tubulin
Real
Naive
Guided
Local Pixel-PCC
-1
1

**Supplementary Figure 2. (A)** Sample segmentation of GM130 labelling (left) using Otsu thresholding (right). **(B)** Sample cell segmentation using Cellpose[32] based on β-catenin labelling. **(C)** Per cell comparison of naïve *versus* guided SemaPlex performance quantified through the Jaccard index calculated between unmixed and real GM130 segmentation. Error bars/box plots in all panels show lower and upper quartiles for 5-fold cross-validated experiments. Significance tested using Welsh's two-sided t-test. **(D)** Comparison of naïve *versus* guided SemaPlex-derived Jaccard indexes for GM130 segmentation per cell. Solid red line shows where the Jaccard index is equivalent for naïve and guided approaches, and dashed lines show ± 0.1 Jaccard index between the two approaches. **(E)** Example cell as presented in Figure 5E showing naïve or guided reconstructions of GM130. The nucleus is denoted by the faded blue DNA overlay. **(F)** Example field of CoxIV, α-tubulin and computational merge of CoxIV and α-tubulin. The DIC image used as the semantic guide in this example is also shown. **(G)** Aggregate comparison of naïve *versus* guided SemaPlex performance quantified as Pixel-PCC calculated between unmixed and real CoxIV (yellow) or unmixed and real α-tubulin (purple). **(H)** Comparison of pixel-PCC scores achieved by naïve *versus* DIC-guided SemaPlex for CoxIV (left) and α-tubulin (right) reconstructions per cell. Solid red line shows where the Jaccard index is equivalent for naïve and guided approaches, and dashed lines show ± 0.01 Jaccard index between the two approaches. **(I)** Visual example of CoxIV and α-tubulin reconstruction. Colour denotes local pixel-PCC score calculated per pixel in a local 3x3 pixel kernel, comparing reconstructed (naïve or guided respectively) *versus* real experimental labelling.
